# PPARδ activation in microglia drives a transcriptional response that primes phagocytic function while countering inflammatory activation

**DOI:** 10.64898/2025.12.17.692628

**Authors:** Jacob S. Deyell, Jonathan Hasselmann, Linda Stroud, Urmimala Raychaudhuri, Yunzi Guo, Byeonggu Cha, Christopher Karma, Deavin Tendean, Minh-Huy Vu Tran, Anastasia Gromova, Audrey S. Dickey, Mathew Blurton-Jones, Albert R. La Spada

## Abstract

Microglia have been implicated in neurodegeneration, though their role remains unclear, as microglia can perform protective functions or promote neuroinflammation. Numerous studies have found that the transcriptome state of microglia can indicate where they lie along this continuum. To understand regulation of microglia transcriptome state, we considered the transcription factor PPAR8, because it is highly expressed in microglia and is a therapeutic target for Alzheimer’s disease, (AD) a neurodegenerative disorder characterized by progressive memory loss where microglia dysfunction is involved. When we delineated the microglia transcriptome in mice treated with PPAR8 agonist, we noted that PPAR8 activation blunted expression of inflammatory mediators and migration-enhancing genes, while boosting phagocytic genes. We then examined PPARδ function in induced transcription factor (iTF) microglia-like cells, and confirmed PPAR8 agonism increases phagocyte function while reducing pro-inflammatory cytokines and migration. To understand PPARδ regulation upon CNS insult, we exposed iTF-microglia to apoptotic neuron debris and defined six microglia transcriptome states as a function of PPAR8 activation, and observed PPAR8 agonism can shift microglia out of a homeostatic state to a primed, disease-associated microglia-like state. As PPAR8 agonism opposed gene expression favored by PU.1, a critical transcription factor in microglial inflammation and AD pathogenesis, we examined their relationship, documented a physical interaction, and found evidence for transrepression. Finally, we tested PPAR8 agonism in Huntington’s disease and tauopathy mice, and demonstrated PPARδ could decrease neuroinflammation in vivo. These findings suggest that PPAR8 agonist therapy may mitigate microglial dysfunction by restoring beneficial functions, while suppressing detrimental inflammation.

## INTRODUCTION

Microglia are the resident immune cells of the central nervous system (CNS)^1^ Under physiological conditions, their main function is surveilling the brain for any foreign material, pathogens, or other “danger signals”. Upon encountering such signals, microglia mount a response, carrying out functions such as chemotaxis, phagocytosis, and cytokine secretion ^1^. Microglia play a central role in neuroinflammation, but their role in neurodegenerative disease is controversial, because while there is evidence for microglial neuroprotection, microglia can also contribute to disease progression ^1,2^. Microglia have been described as “activated” in the setting of neurodegenerative disease, and then to take on distinct disease-related phenotypes; however, the activation state of microglia varies based upon their environmental situation, and thus their characterization in different contexts, including that of neurodegenerative disease, requires assessment of both gene expression and function ^3^. One of the first neurodegeneration-associated transcriptomic states of microglia to be delineated was the TREM2-dependent Disease Associated Microglia (DAM) state or microglia neurodegenerative (MGnD) phenotype, which may reflect a protective compensatory state characterized by a downregulation of homeostatic markers (e.g., *P2RY12* and *TMEM119*) and upregulation of pro-inflammatory and phagocytic capacity genes (e.g., *CLEC7A* and *SPP1*) ^4,5^. However, recent single-cell experiments have demonstrated that microglia states in neurodegeneration are very diverse, with one dataset demonstrating 12 different microglia states in the human Alzheimer’s disease brain ^6^.

Alzheimer’s disease (AD) is a devastating neurodegenerative disorder characterized by progressive memory loss and impairment of behavioral functions ^7^. Neuropathologically, it is classified by the accumulation of amyloid-beta (Aβ) plaques and neurofibrillary tau tangles in the brain ^7^. These protein aggregates lead to neuronal dysfunction and death, resulting in a gradual decline in cognitive abilities. AD is the most common cause of cognitive decline, and sporadic AD accounts for ∼90% of all cases^8^. While *APOE* allele status is a predominant genetic risk factor for AD ^8^, over the last two decades, unbiased genome-wide association studies (GWAS) of sporadic AD have identified a number of risk loci encoding genes that are expressed primarily or exclusively in microglia ^1^, implicating dysfunction of microglia as a likely shared pathway of AD pathogenesis. Though there is no known single cause of sporadic AD, certain genetic risk factors predispose individuals to developing the disease. The *APOE4* allele is the single greatest genetic risk factor for AD^8^. Microglia harboring the *APOE4* allele have been extensively characterized and have been shown to display deficient phagocytic capacity, secrete excessive inflammatory cytokines, and be unable to adopt the DAM state ^9,10^. They have also more recently been described to exhibit severe lipid dysregulation, leading to the accumulation of lipid droplets ^11^. Another well-studied AD risk factor is PU.1, a master transcription factor encoded by the *Spi1* gene ^12^. PU.1, which is crucial for regulation of homeostasis and inflammatory responses, is an established AD risk factor and controls the expression of several other AD genetic risk factors, including *TREM2* and *APOE* ^12^. Consequently, considerable effort has been focused on identifying modulators of PU.1 activity, with a recent study demonstrating neuroprotection in AD models when PU.1 activity is decreased ^13^.

While understanding the role of AD genetic risk factors is crucial, exploring therapeutic targets that can modulate microglia behavior has emerged as a key focus in AD translational research. One such potential target is peroxisome proliferator-activated receptor delta (PPARδ), a member of the PPAR family, which consists of two other members - PPARα and PPARγ. The PPARs are ligand-activated transcription factors in the nuclear hormone receptor superfamily, initially studied in the context of lipid metabolism and metabolic disease due to their role as lipid sensors ^14^. Beyond this action, however, the PPARs are crucial regulators of inflammation ^15^. PPAR8 is highly expressed in the CNS, and we previously documented that PPARδ dysregulation in neurons accurately recapitulates Huntington’s disease (HD) phenotypes in mice, and found that PPAR8 agonist treatment could rescue HD neurodegeneration in model mice and patient-derived striatal neurons ^16^. Interestingly, microglia are among the highest expressors of PPARδ in the CNS, and PPAR8 is the most highly expressed PPAR in microglia ^17,18^. The PPAR8 agonist T3D-959 has been studied as a potential treatment for sporadic AD, and has shown some promise in human patient clinical trials ^19^. Microglial PPAR8 is also neuroprotective in models of Experimental Autoimmune Encephalomyelitis (EAE), where it has been shown to exhibit anti-inflammatory effects ^20,21^, but exactly how PPAR8 regulates microglia function is yet to be determined.

Here we performed unbiased RNA-seq transcriptome analysis to evaluate microglia in mice subjected to PPAR8 agonism *in vivo*, and documented a blunting of microglial inflammation upon PPAR8 activation. To better comprehend the action of PPARδ in microglia, we pursued a combined analysis of transcriptional state and cellular function in induced microglia-like cells (iMGs) derived from induced pluripotent stem cells (iPSCs) through the use of ectopically expressed transcription factors ^22^. Our studies of these induced transcription factor (iTF)-microglia revealed that PPAR8 agonist treatment shifted microglia from baseline homeostasis to a phagocytic primed state, with increased lipid mobilization and processing capacity, but without pronounced inflammation. Because our findings revealed that PPARδ counters the action of PU.1 in microglia, we explored the relationship between PPARδ and PU.1, and found that they physically interact and promote opposing microglial functions. We then evaluated the effect of PPAR8 agonist treatment on neuroinflammation occurring in the context of neurodegenerative disease in HD and tauopathy model mice, and noted marked reductions in the expression of NF-kB-dependent inflammatory cytokine expression in the brains of PPAR8-treated animals. Our results indicate that PPAR8 activation in microglia can reset their expression and functional state to favor phagocytic capacity while blunting activation of inflammatory pathways, and thus nominate PPAR8 agonist therapy as a microglia-modifying treatment for AD and related neurodegenerative disorders.

## RESULTS

### PPAR8 agonism reduces inflammation in microglia *in vivo*

To determine the effect of PPAR8 activation on microglia, we began by testing if delivery of a brain-penetrant PPAR8 agonist would modulate the microglia transcriptome in wild-type adult C57BL/6J mice. After a six-week treatment period during which mice received either the PPAR8 agonist KD3010 or vehicle control, we isolated microglia and performed bulk RNA sequencing. This transcriptome analysis yielded 1,947 differentially expressed genes (DEGs), consisting of 1,016 significantly down-regulated genes and 931 significantly up-regulated genes (*P*_adj_ < 0.05) (**Fig. 1a**). To validate these findings, we confirmed a subset of hits by qRT-PCR analysis (**Fig. 1b**). We then performed gene ontology (GO) analysis, which revealed strong evidence for reduced immune activation, based upon evaluation of significantly down-regulated DEGs (**Fig. 1c**). As PPARs are known to exhibit prominent anti-inflammatory effects in addition to their well-established role in regulating lipid metabolism ^15,23^, our results indicate that PPAR8 activation in microglia may promote reduced inflammation in the brain. Additionally, cell migration was a top hit of downregulation by PPAR8 agonism. Migration of microglia is intimately tied to their immune response; thus, decreased migration may blunt the extent of inflammation in response to a stimulus. We also evaluated the biological implications of the significantly up-regulated DEGs detected in the microglia of adult WT mice treated with PPAR8 agonist by performing KEGG analysis, and we noted that the “endocytosis” pathway was highly significant (**Supplementary Table 1**), suggesting that phagocytosis might be enhanced in microglia subjected to PPAR8 activation.

**Figure 1.**
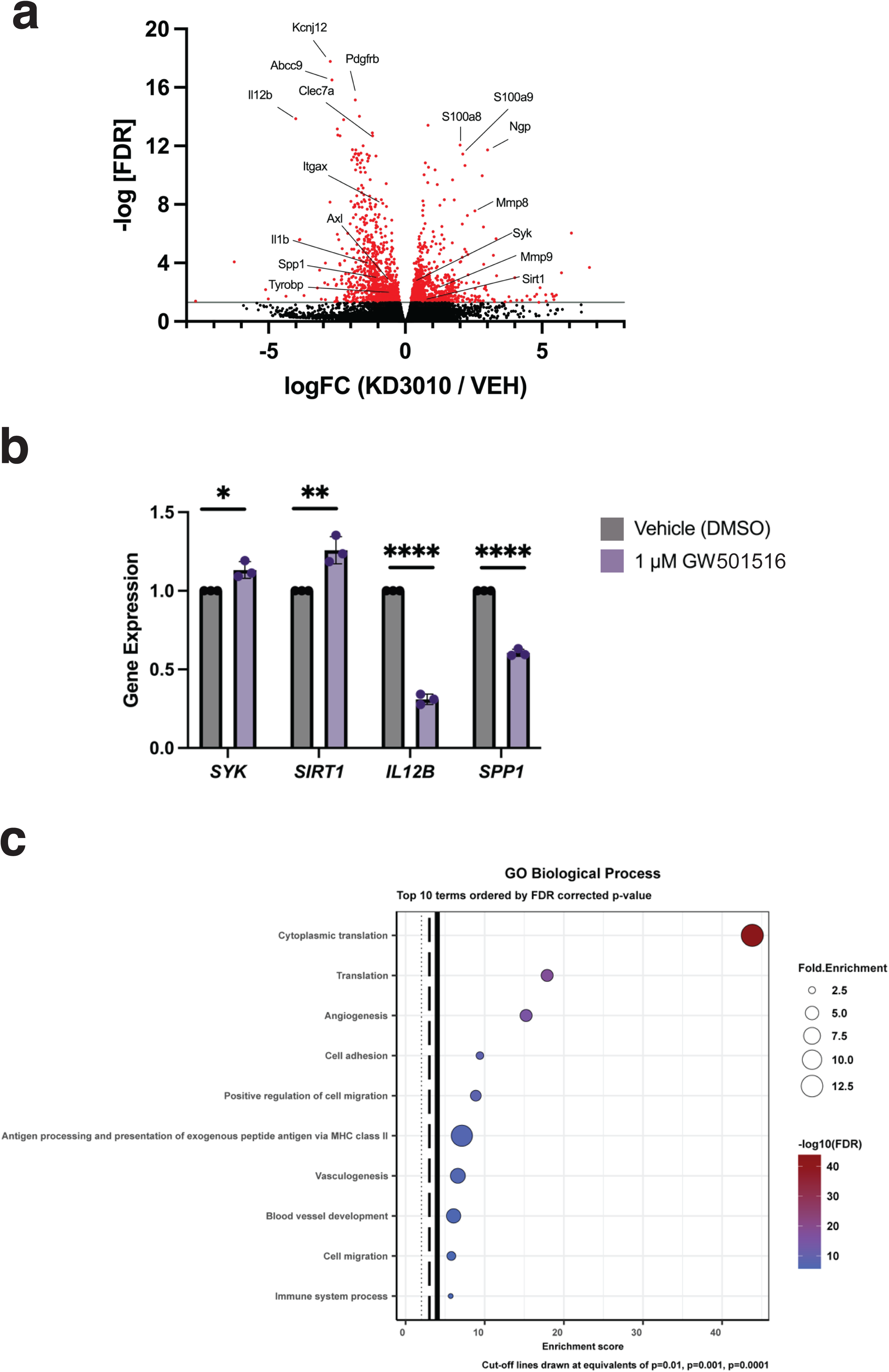
Bulk RNA-seq analysis of mice treated with PPAR8 agonist. **a)** Volcano plot showing up- and down-regulated genes of microglia in wild-type C57BL/6J mice upon PPAR8 agonist treatment, with select genes labeled (n = 4 mice/group). Significance level of p_adj<0.05. **b)** We performed qRT-PCR validation of select PPAR8 target genes in iTF-Microglia. *p<0.05; **p<0.01; ****p<0.0001, two-tailed t-test, n = 3 biological replicates. **c)** Gene ontology analysis of down-regulated microglia biological processes resulting from PPAR8 agonist *in vivo* treatment of wild-type C57BL/6J mice. Categories are sorted by -log10(FDR) with vertical cut-off lines drawn at certain adjusted p-values (dotted: p=0.01, dashed: p=0.001, solid: p=0.0001). Error bars = s.e.m.

Expression changes in certain single genes with important regulatory roles were also noted in mouse brain microglia upon PPAR8 agonist activation. In response to PPAR8 agonism, microglia upregulated expression of *Syk* and *Sirt1*, while reducing expression of *IL12b* and *Spp1*. SYK is a key regulator of phagocytosis and enhances a neuroprotective microglia response in neurodegenerative disease, and its deletion exacerbates neuropathology and cognitive impairment in AD model mice ^24^. SIRT1, a member of the well-studied sirtuin family, reduces neuroinflammation and is also neuroprotective in AD ^25,26^. Thus, upregulation of these genes could provide benefit in the setting of AD. Conversely, IL12b is a subunit of the inflammatory cytokines IL-12 and IL-23, which are upregulated in response to amyloid beta (Aβ) and whose inhibition attenuates AD pathology ^27^. SPP1 (osteopontin) is a cytokine shown to impair Aβ uptake, suggesting that its downregulation may also be neuroprotective by priming microglia for sustained phagocytic demand ^28^. These initial *in vivo* findings thus demonstrate that PPAR8 agonism in microglia may prevent pathological inflammation while promoting phagocytic capacity.

### PPARδ reduces pro-inflammatory cytokine release from microglia and blunts migration

To directly evaluate the effect of PPAR8 activation on microglia, we utilized a system for deriving microglia from iPSCs by applying a direct cell fate conversion strategy predicated on induction of six key transcription factors, resulting in production of highly-enriched microglia-like cells, known as iTF-microglia ^22^. To achieve transition to an inflammatory state, we treated iTF-microglia with lipopolysaccharide (LPS), a bacterial endotoxin widely used as a potent inducer of inflammatory pathways through the NF-kB pathway and Toll-Like Receptor 4 pathway ^29^, and we noted a robust inflammatory response, based upon detection of elevated levels of 12 inflammatory cytokines (**Fig. 2a, b**), in agreement with previously published studies ^22^. However, when we co-treated iTF-microglia with LPS and the PPAR8 agonist GW501516, we observed a marked reduction in inflammatory cytokine release and noted an increase in the level of the anti-inflammatory cytokine IL-10 (**Fig. 2a, b**).

**Figure 2.**
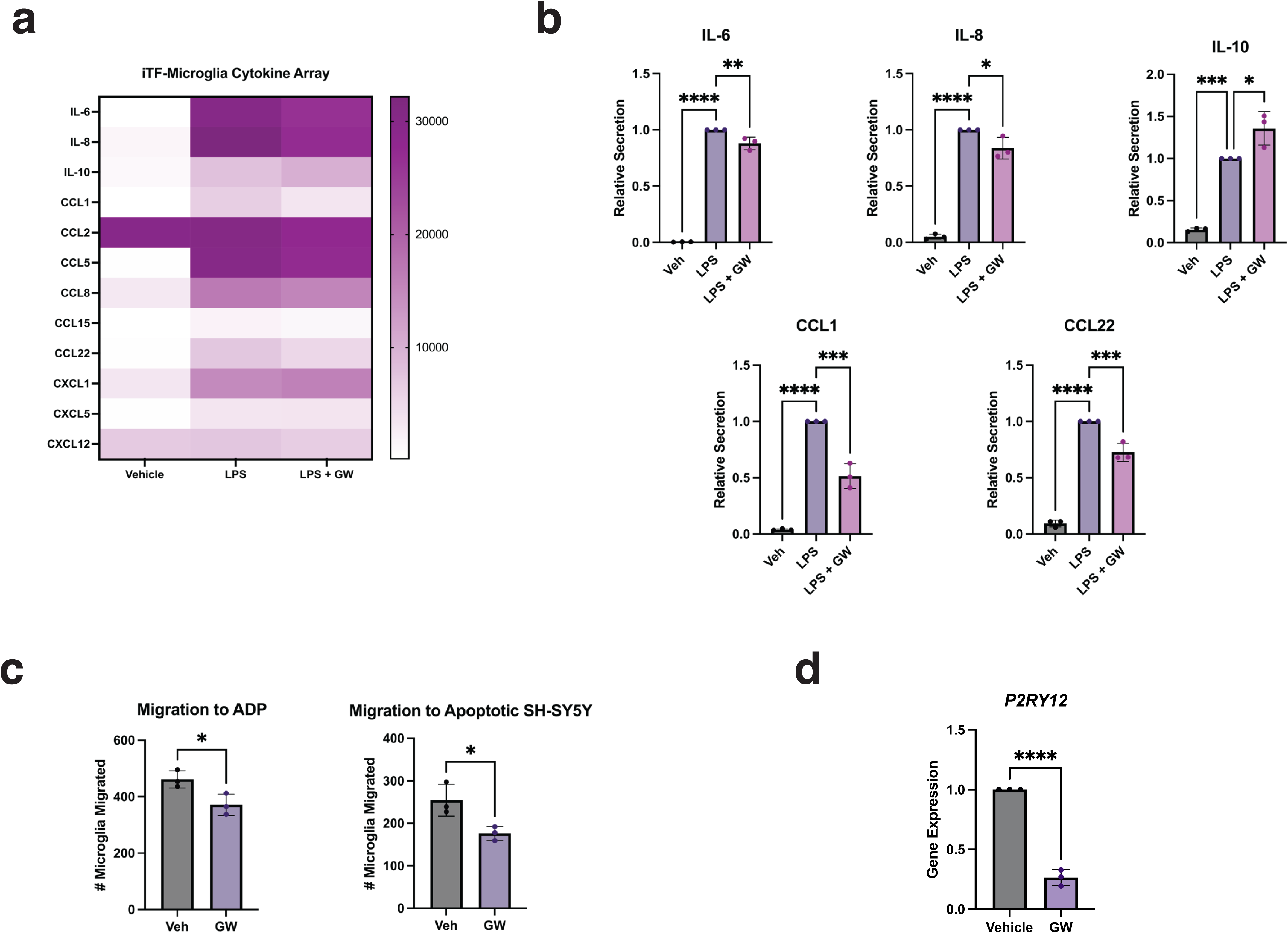
PPAR8 agonismt treatment reduces inflammation in LPS-stimulated iTF-microglia. **a)** iTF-microglia were stimulated with LPS alone or in combination with GW501516 treatment. Conditioned media from the cells was removed and run on a cytokine array to measure quantities of secreted cytokines, with the heatmap showing cytokines that were significantly altered by LPS and how these were changed by PPAR8 agonist treatment. Average raw readings plotted, n = 3 biological replicates. **b)** Quantification of results from (A) for cytokines showing significant changes upon PPAR8 agonist treatment of LPS-stimulated iTF-microglia. *p<0.05; **p<0.01; ***p<0.001; ****p<0.0001, one-way ANOVA with post-hoc Tukey test, n = 3 biological replicates. **c)** iTF-microglia treated with the PPAR8 agonist GW501516 were incubated with two different stressors, and numbers of migrating microglia were counted. Phagocytosis (fluorescence) was monitored over 24 hours (n = 4 independent wells; 4 images per well). Images were acquired every 30 minutes using the IncuCyte S3 live imaging system. *p<0.05, two-tailed t-test, n = 3 biological replicates. **d)** We performed qRT-PCR analysis of *P2RY12* expression level in iTF-microglia subjected to 24 hours treatment with the PPAR8 agonist GW501516 (1 μM). ****p<0.0001, two-tailed t-test, n = 3 biological replicates. Error bars = s.e.m.

Another key function of microglia as immune cells is migration, which is involved in the microglial response to injury. As PPAR8 activation in endogenous mouse microglia resulted in downregulation of genes associated with the “Cell migration” and “Positive regulation of cell migration” pathways (**Fig. 1c**), we chose to next examine migration via a transwell assay system. To accomplish this, iTF-microglia were plated onto a removable mesh insert placed into a well that contained an attractant and the number of microglia moving through the pore were quantified as a read-out of migration (**Supplementary Fig. 1**). As migration assays have not yet been routinely conducted on iTF-microglia, we tested a variety of potential attractants and found that ADP and apoptotic SH-SY5Y cells were potent inducers of migration. To assay migration of PPAR8-activated iTF-microglia, we treated iTF-microglia with the PPAR8 agonist GW501516 or with vehicle for one hour, and then exposed the iTF-microglia to attractant in the transwell system. We found that pre-treatment with PPAR8 agonist significantly reduced the extent of iTF-microglia migration (**Fig. 2c**). Both ADP and apoptotic SH-SY5Y cells (which release ADP) act on microglia via the purinergic receptor P2RY12, a well-known marker of microglial homeostasis ^30^. To determine if PPARδ directly affects P2RY12 expression in microglia, we again treated iTF-microglia with the PPAR8 agonist GW501516 or with vehicle for one hour, measured the level of *P2RY12* via qRT-PCR analysis, and documented a significant reduction in *P2RY12* expression (**Fig. 2d**). Hence, PPAR8 may blunt microglia migration by repressing *P2RY12* expression.

### PPAR8 activation increases microglia phagocytosis of AD-relevant substrates

One important neuroprotective function of microglia is to remove pathological accumulation of misfolded proteins and eliminate cellular debris from the CNS. Indeed, genome-wide association studies of AD have identified risk alleles in certain microglia expressed genes, including *TREM2* and *APOE*, and previous studies have documented that mutant versions of these proteins, when expressed in microglia, can interfere with the normal homeostatic function of microglia, thereby predisposing to AD pathogenesis ^9,10,31,32^. To determine if PPAR8 activation modulates phagocytic function, we treated iTF-microglia with the PPAR8 agonist GW501516 or vehicle, and we incubated iTF-microglia with either fluorescent fibrillar Aβ_1-42_ or pHrodo red-labelled rat synaptosomes. We tracked uptake of fluorescent fibrillar Aβ_1-42_ by iTF-microglia over the course of 24 hours, and we found that PPAR8 activation of iTF-microglia enhanced uptake of fAβ_1-42_ (**Fig. 3a**). Although microglial phagocytic function against fAβ_1-42_ roughly doubled at the 24 hour mark in iTF-microglia treated with PPAR8 agonist, this only constituted a strong trend (**Fig. 3b**). However, when we tracked uptake of fluorescent pHrodo red-labelled rat synaptosomes, we detected a dramatic increase in phagocytosis by iTF-microglia treated with PPARδ (**Fig. 3c**). In this case, increased phagocytosis elicited by PPAR8 agonist treatment was significantly greater than the phagocytosis occurring in vehicle-treated iTF-microglia (**Fig. 3d**). These findings indicate that activation of PPARδ can boost microglia phagocytic capacity.

**Figure 3.**
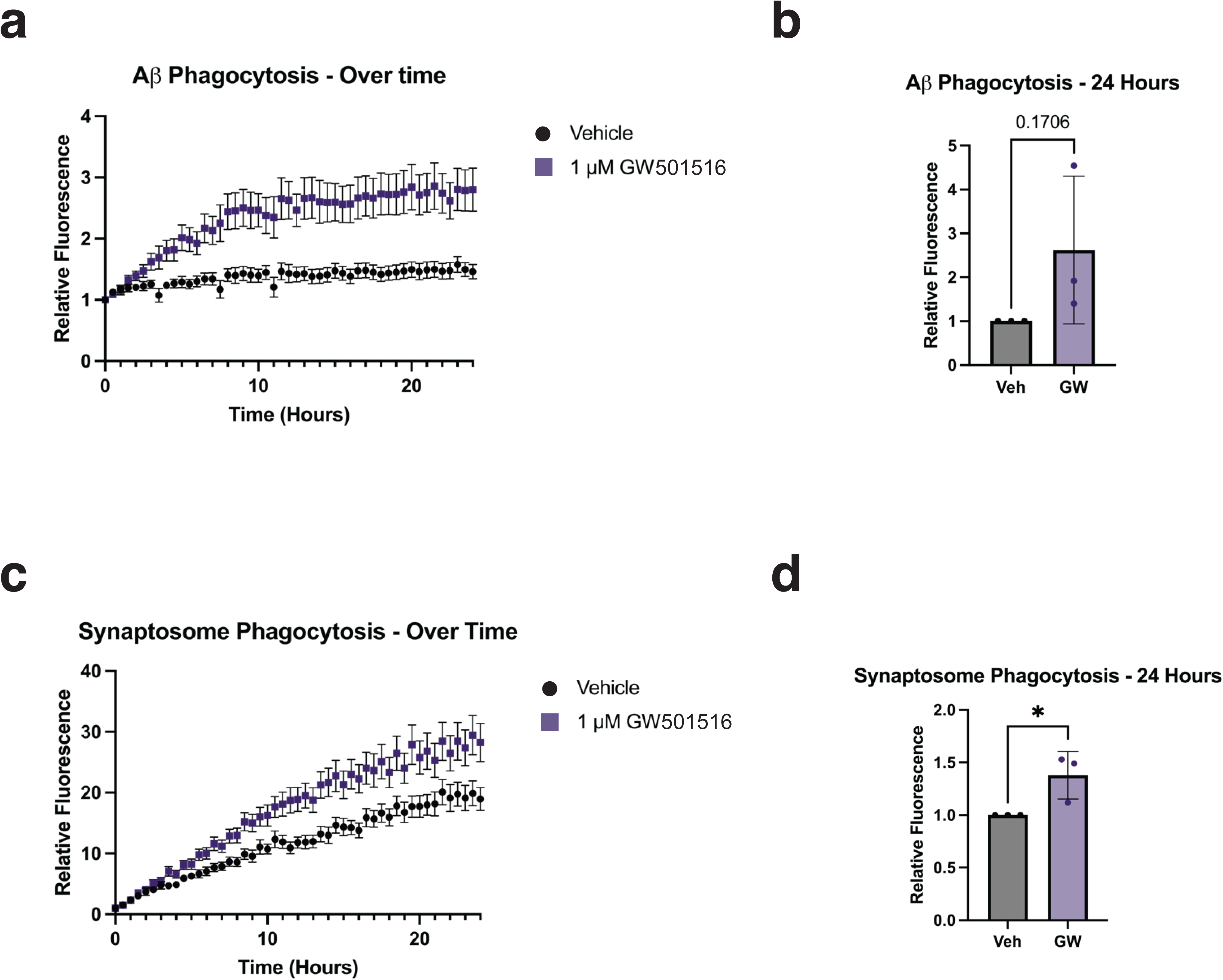
PPAR8 agonism upregulates the phagocytosis of disease-relevant substrates. **a)** iTF-Microglia were incubated with fibrillar Aβ (5 μg/mL), with or without GW (1 μM). Phagocytosis (fluorescence) was monitored over 24 hours (n = 4 independent wells; 4 images per well). Images were acquired every 30 minutes using the IncuCyte S3 live imaging system. n = 3 biological replicates. **b)** Quantification of fibrillar Aβ phagocytosis at the 24-hour timepoint. Two-tailed t-test, n = 3 biological replicates. **c)** iTF-Microglia were incubated with rat synaptosomes (1 mg/mL) with or without GW (1 μM). Phagocytosis (fluorescence) was monitored over 24 hours (n = 4 independent wells; 4 images per well). Images were acquired every 30 minutes using the IncuCyte S3 live imaging system. n = 3 biological replicates. **d)** Quantification of synaptosome phagocytosis at the 24-hour timepoint. *p < 0.05, two-tailed t-test, n = 3 biological replicates. Error bars = s.e.m.

### iTF-microglia respond to PPARδ modulation and broad neurodegenerative challenge by altering lipid processing, inflammation, and cell migration genes

As our results demonstrate that PPARδ agonism enhanced amyloid phagocytosis while reducing LPS-induced inflammation, we next sought to assess how PPARδ activity would affect the microglial response to a broader CNS insult. Numerous studies have documented how microglia function and how gene expression changes in the setting of neurodegenerative disease, including especially AD ^3^. It has also been reported previously that a DAM-like state can be induced by treating iPSC-derived microglia with myelin debris or apoptotic neurons ^33^, and we utilized a novel, more comprehensive insult involving the use of “brain powder” prepared by flash freezing aged mouse brains and pulverizing to a fine powder. To assess how PPARδ activation can modulate microglial state, we exposed DIV9 iTF-microglia to saline or brain powder in combination with PPARδ agonist or PPARδ antagonist drug treatment, and then performed Parse single cell RNA sequencing (scRNA-seq).

Following quality control to remove low quality cells (**Supplementary Fig. 2**), we evaluated the iTF-microglia transcriptome response to PPARδ agonist or PPARδ antagonist in the absence of brain powder (**Supplementary Table 2**). This analysis enabled us to map 16,916 cells across the three treatment groups to six unique clusters (**Fig. 4a**; **Supplementary Table 3**). Comparison of the cell cluster distributions revealed many significant alterations, with the greatest changes observed in clusters Mg1, Mg4, and Mg5. As expected, treatment with PPARδ agonist yielded cell cluster shifts directly opposite to those observed upon treatment with PPARδ antagonist, confirming that these drugs elicit contrary effects on microglia function (**Fig. 4b**). To validate these findings, we measured the expression of *PDK4* and *ANGPTL4*, two canonical PPARδ target genes, and found that both genes were significantly upregulated upon PPARδ activation, but significantly downregulated with PPARδ inhibition (**Fig. 4c**). We next evaluated genes typically expressed in homestatic microglia, including *ABCA1*, a transporter that moves cholesterol out of the cell; *CYP1B1*, a lipid metabolism enzyme; and *PLIN2*, a key component of intracellular lipid droplets, and we documented that expression of all three genes are significantly upregulated upon PPARδ agonist treatment, while expression of *ABCA1* and *PLIN2* are significantly decreased upon PPARδ antagonist treatment (**Fig. 4d**). When we measured expression levels of the inflammatory cytokine genes *CCL13*, *CCL2*, and *IL1B*, we observed marked decreases with PPAR8 agonism (**Fig. 4e**); however, PPARδ antagonism did not affect the expression of these genes, as exposure to an inflammatory stimulus might be required to elicit such an effect. Finally, when we evaluated the ADP-sensing receptors *P2RY12* and *P2RY13*, and the lipid-sensing scavenger receptor *TREM2*, which has been shown to play a critical role in microglia migration towards amyloid-beta plaques via its association with TYROBP ^4,31^, we documented significant coordinate downregulation of all three genes following PPARδ agonism, and detected significant expression increases for *P2RY12* and *P2RY13* with PPARδ antagonism (**Fig. 4f**).

**Figure 4.**
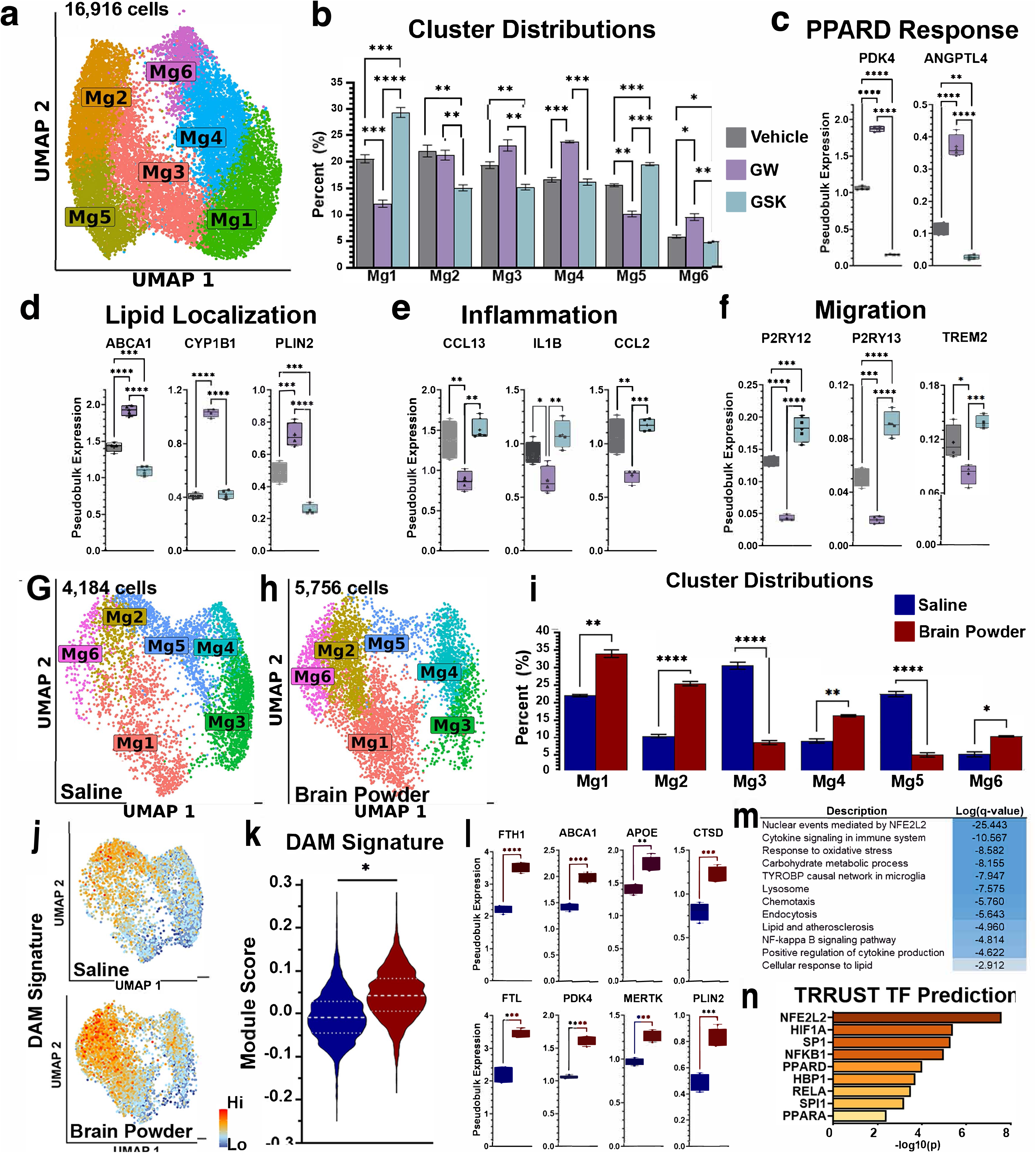
iTF-microglia response to PPARd modulation and brain powder stimulation. **a)** UMAP showing the clustering of iTF-Microglia treated with vehicle, GW, or GSK. **b)** Barplot detailing the cluster distributions of each treatment group within the UMAP in ’A’. *p<0.05; **p<0.01; ***p<0.001; ****p<0.0001, two-way ANOVA with Tukey’s test for multiple comparisons. **c)** Boxplots detailing the normalized pseudobulk expression levels for the PPAR8 responsive genes *PDK4* and *ANGPTL4*. **p<0.01; ****p<0.0001, one-way ANOVA with post-hoc Tukey test. **d)** Boxplots detailing the normalized pseudobulk expression levels for the lipid localization genes *ABCA1*, *CYP1B1*, and *PLIN2*. ***p<0.001; ****p<0.0001, one-way ANOVA with post-hoc Tukey test. **e)** Boxplots detailing the normalized pseudobulk expression levels for the inflammation associated genes *CCL13*, *IL1B*, and *CCL2*. *p<0.05; **p<0.01; ***p<0.001, one-way ANOVA with post-hoc Tukey test. **f)** Boxplots detailing the normalized pseudobulk expression levels for the migration associated genes *P2RY12*, *P2RY13*, and *TREM2*. *p<0.05; ***p<0.001; ****p<0.0001, one-way ANOVA with post-hoc Tukey test. **g-h)** UMAPs detailing the cluster distributions of iTF-microglia treated with saline (G) or brain powder (H) after data integration and clustering as a merged dataset. **i)** Barplot detailing the cluster distributions of each treatment group within the UMAP in ’G-H’. *p<0.05; **p<0.01; ****p<0.0001, two-way ANOVA with Tukey’s test for multiple comparisons. **j)** Heatmaps showing the intensity and localization of the DAM gene signature in iTF-Microglia treated with saline or brain powder. **k)** Violin plot detailing the module scores for the DAM gene signature in all cells. *p<0.05, two-tailed t-test. **l)** Boxplots detailing the normalized pseudobulk expression levels for multiple significantly upregulated DAM signature genes. **p<0.01; ****p<0.0001, two-tailed t-test. **m)** A subset of pathways identified by performing Metascape enrichment analysis on genes that are upregulated (FDR<0.05; LFC>0.25) in brain powder-treated iTF-microglia vs saline-treated iTF-microglia. **n)** Transcription factors identified via Metascape’s query of the TRRUST database as being significantly related to the gene set that is upregulated following powder treatment. For all boxplots, bars denote the minimum and maximum points, the central line denotes the median, and the “+” denotes the mean. Error bars = s.e.m.

After determining that iTF-microglia respond to PPARδ agonism and antagonism in a diametrically opposed fashion, we next sought to verify that the brain powder treatment induced an activation state resembling the *in vivo* microglia response that occurs in the vicinity of amyloid-beta plaque pathology. Brain powder-treated samples were clustered to reveal 4,184 saline treated cells and 5,756 brain powder treated cells that separated into six distinct cell clusters (**Fig. 4g, h**). Brain powder treated cells were significantly enriched in clusters Mg1, Mg2, Mg4, and Mg5 (**Fig. 4i**), with clusters Mg2 and Mg5 showing the highest levels of gene expression associated with the DAM signature (**Fig. 4j**; **Supplementary Table 3**). The DAM gene expression signature was significantly upregulated across all brain powder treated cells (**Fig. 4k**), with notable expression increases for DAM genes involved in lipid processing, lysosomal function, and iron homeostasis (**Fig. 4l**). When we performed gene annotation analysis using Metascape^34^, we noted several significantly upregulated pathways related to cytokine signaling, TYROBP signaling, lysosome function, endocytosis, and cellular lipid responses in the brain powder treated cells (**Fig. 4m**). Furthermore, the genes driving these pathway predictions were significantly associated with a number of transcription factors, including PPARδ, PPARα, PU.1, NFE2L2 and NFKB1 (**Fig. 4n**).

### Microglial PPARδ modulation regulates lipid homeostasis, mitochondrial fatty acid beta-oxidation, and inflammation in response to neurodegenerative insult

As our transcriptome data indicate that PPARδ modulation affects microglia gene expression and brain powder treatment can successfully activate microglia, based upon gene expression, pathway, and transcription factor alterations, we next sought to determine how PPARδ regulates the microglial response to CNS injury. Brain powder treated samples were separated from the Seurat object and reclustered together to reveal 16,829 cells distributed across seven clusters (**Fig. 5a, b**). Examination of cluster distributions across the three treatment groups revealed several significant shifts, although only cluster Mg3 showed opposing cluster shifts in PPARδ agonist vs. PPARδ antagonist samples (**Fig. 5c**). To determine which of these phenotypic alterations reflected PPARδ target gene activation, we performed pathway analysis on significantly upregulated DEGs in both PPARδ agonist and PPARδ antagonist conditions (**Fig. 5d-i**). Genes upregulated upon PPARδ antagonism revealed a significant association with pathways related to the inflammatory response, highlighting transcription factors encoded by the *RELA*, *NFKB1*, and *SPI1* genes (**Fig. 5d, e; Supplementary Table 4**). The aggregated expression of a subset of genes from the “Inflammatory Response” pathway revealed that these genes are most highly enriched in cluster Mg3, a cluster that is significantly reduced with PPARδ agonism and increased with PPARδ antagonism, confirming that PPARδ activation can blunt microglial inflammation (**Fig. 5f**; **Supplementary Table 4**). Conversely, genes upregulated upon PPARδ agonism are associated with lipid metabolism, fatty acid beta oxidation, PPAR signaling, and endocytosis pathways (**Fig. 5g**).

**Figure 5.**
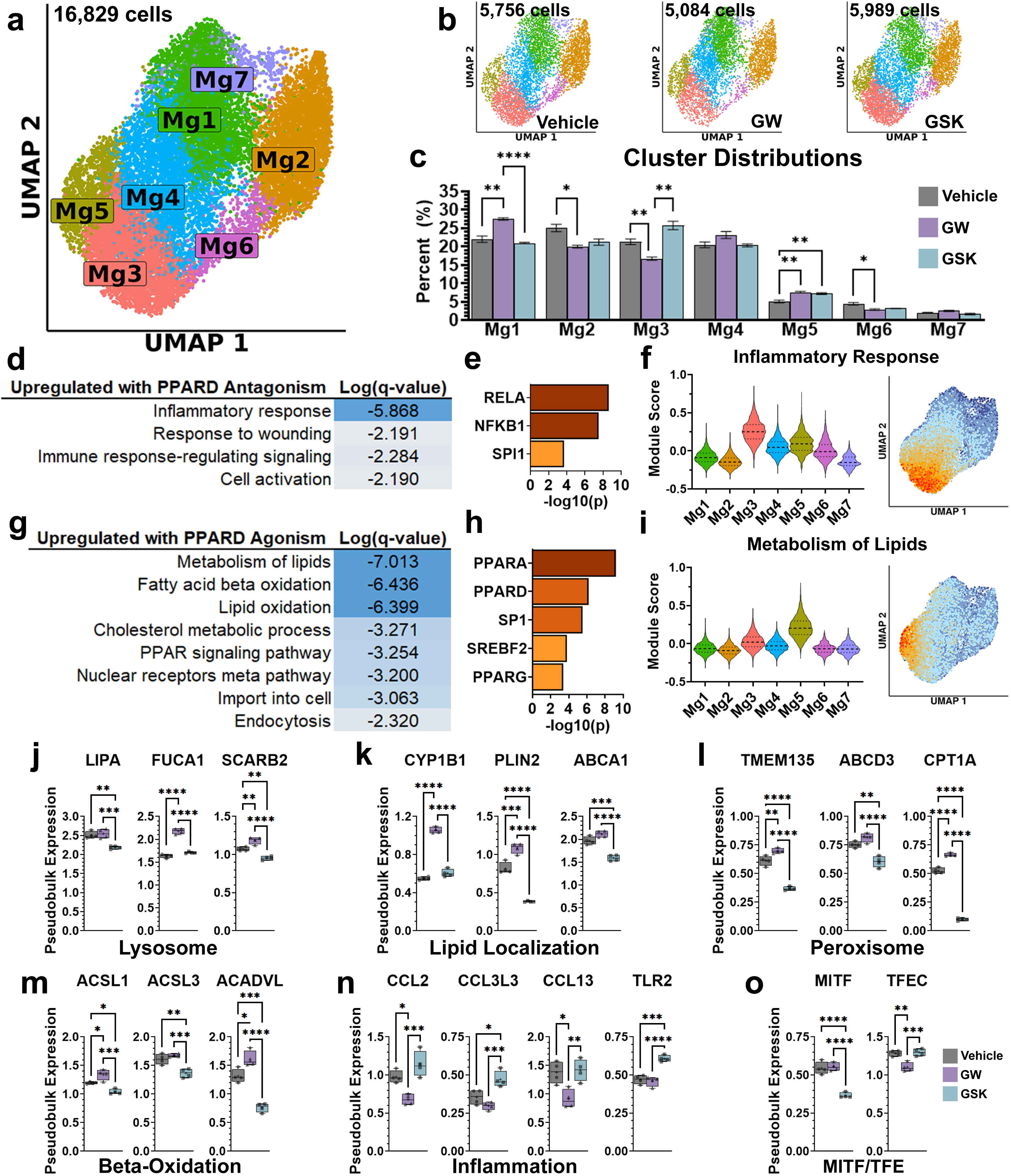
Effect of PPAR8 modulation on iTF-microglia response to brain powder treatment. **a)** UMAP showing the clustering of brain powder-treated iTF-microglia that also received vehicle, GW501516, or GSK3787 treatment. n = 4 biological replicates. **b)** Split UMAPs showing cluster distributions of cells from each treatment group. **c)** Barplot detailing the cluster distributions of each treatment group within the UMAPs in ’A-B’. *p<0.05; **p<0.01; ****p<0.0001, two-way ANOVA with Tukey’s test for multiple comparisons. **d)** Pathways identified by performing Metascape enrichment analysis on genes that are downregulated (FDR<0.05; LFC<-0.25) in GW5051516-treated iTF-microglia vs GSK3787-treated iTF-microglia. **e)** Transcription factors identified via Metascape’s query of the TRRUST database as being significantly related to the gene set in ’D’. **f)** Violin plots detailing the module scores for a subset of genes in the GO Biological Processes “Inflammatory Response” pathway across all cells in each cluster. Dashed lines represent the median (center) and quartiles (upper and lower). **g)** Pathways identified by performing Metascape enrichment analysis on genes that are upregulated (FDR<0.05; LFC>0.25) in GW501516-treated iTF-Microglia vs GSK3787-treated iTF-microglia. **h)** Transcription factors identified via Metascape’s query of the TRRUST database as being significantly related to the gene set in ’G’. **i)** Violin plots detailing the module scores for a subset of genes in the Reactome “Metabolism of Lipids” pathway across all cells in each cluster. Dashed lines represent the median (center) and quartiles (upper and lower). **j)** Boxplots detailing the normalized pseudobulk expression levels for lysosome genes *LIPA*, *FUCA1*, and *SCARB2*. **p<0.01; ***p<0.001; ****p<0.0001, one-way ANOVA with post-hoc Tukey test. **k)** Boxplots detailing the normalized pseudobulk expression levels for lipid localization genes *CYP1B1*, *PLIN2*, and *ABCA1*. ***p<0.001; ****p<0.0001, one-way ANOVA with post-hoc Tukey test. **l)** Boxplots detailing the normalized pseudobulk expression levels for peroxisome genes *TMEM135*, *ABCD3*, and *CPT1A*. **p<0.01; ****p<0.0001, one-way ANOVA with post-hoc Tukey test. **m)** Boxplots detailing the normalized pseudobulk expression levels for mitochondrial fatty acid beta-oxidation genes *ACSL1*, *ACSL3*, and *ACADVL*. *p<0.05, **p<0.01; ***p<0.001; ****p<0.0001, one-way ANOVA with post-hoc Tukey test. **n)** Boxplots detailing the normalized pseudobulk expression levels for inflammation-associated genes *CCL2*, *CCL3L3*, *CCL13*, and *TLR2*. *p<0.05, **p<0.01; ***p<0.001; ****p<0.0001, one-way ANOVA with post-hoc Tukey test. **o)** Boxplots detailing the normalized pseudobulk expression levels for transcription factors *MITF*, and *TFEC*. **p<0.01; ***p<0.001; ****p<0.0001, one-way ANOVA with post-hoc Tukey test. For all boxplots, bars denote the minimum and maximum points, the central line denotes the median, and the “+” denotes the mean. Error bars = s.e.m.

Corrspondingly, these genes are most significantly associated with the three PPAR transcription factors and SREBF2, a regulator of cholesterol homeostasis (**Fig. 5h**). Further examination of a subset of the “Metabolism of Lipids” pathway genes revealed that cells in cluster Mg5 were the most enriched for expression of this lipid processing gene program (**Fig. 5i**). Interestingly, both PPARδ agonism and antagonism significantly increased the proportion of microglia in cluster Mg5, suggesting that promotion of lipid processing is not fully inhibited when PPARδ activity is blocked.

To further dissect how PPARδ is regulating these phenotypic shifts, we sought to more specifically examine how genes across multiple related cellular compartments are altered in response to PPARδ modulation. Given that microglia lipid processing involves endocytosis, lysosome fusion, storage within intracellular lipid droplets, or secretion of excess cholesterol, and that further fusion of lipid droplets with peroxisomes prior to their utilization is an energy source via mitochondrial fatty acid beta-oxidation, we focused on genes associated with these cellular components. The lysosomal genes *LIPA*, *FUCA1*, and *SCARB2* all showed significant downregulation upon PPARδ antagonism, and significant upregulation upon PPARδ agonism, suggesting that PPARδ maintains or enhances lysosome function in microglia (**Fig. 5j**). Turning to genes involved in cellular lipid localization, *CYP1B1*, *PLIN2*, and *ABCA1* yielded mixed effects, but indicated that PPARδ activation can enhance multiple components of lipid processing, while PPARδ inhibition will reduce microglial capacity for moving lipid through the cell (**Fig. 5k**). Importantly, the lipid droplet gene *PLIN2* was significantly increased with PPARδ agonism and significantly decreased with PPARδ antagonism, suggesting that microglial capacity for storing lipids is directly regulated by PPARδ. We next looked at genes associated with the peroxisome and mitochondrial beta-oxidation and again found that the overall trend is that PPARδ agonism enhances these pathways, while antagonism directly inhibits these pathways in microglia (**Fig. 5l, m**).

Focusing next on inflammatory pathways, we examined the expression of *CCL2, CCL3L3, CCL13*, and *TLR2* (**Fig. 5n**). As observed previously, PPARδ agonist-treated cells exhibited significantly decreased levels of these inflammatory genes, while PPARδ antagonist treatment did not greatly increase the expression of these genes. However, these genes are all most highly expressed in cluster Mg3, which is significantly increased upon PPARδ antagonism, but significantly decreased with PPARδ activation (**Fig. 5c**). These findings indicate that although PPARδ activation can suppress inflammatory gene expression, PPARδ inhibition may regulate the inflammatory state by reducing the overall number of cells primed to transition to an inflammatory state.

Finally, while predictive analysis identified gene program alterations that were associated with RELA, NFKB1, PU.1, and the three PPAR transcription factors, we also observed significant expression changes for many other transcription factors (**Supplementary Table 3**). Of particular interest were significant alterations in *MITF* and *TFEC*, two members of the MITF/TFE transcription factor family. Our results indicate that PPARδ antagonism significantly reduced the expression of *MITF*, while PPARδ agonism significantly reduced *TFEC* expression (**Fig. 5o**). These transcription factors are of particular interest, as the MITF/TFE family is well known for regulating lysosome function^35^ and previous work has proposed MITF as a key regulatory node in the microglial DAM phenotype ^33^. As transcription factors in the MITF/TFE family can form homodimers and heterodimers prior to translocating to the nucleus ^35^, our findings suggest that PPARδ may play an upstream role in modulating the concentration of various MITF/TFE family members, thereby influencing which factors dimerize to control downstream microglial gene expression.

### PPARδ interacts with PU.1 and represses its activity

One intriguing result of our studies is that PPAR8 activation can potently blunt the expression of inflammatory genes regulated by the transcription factor PU.1, which is the product of the *SPI1* gene. PU.1 is a master transcription factor in microglia that has been found to regulate the expression of numerous AD risk genes and is itself a genetic risk factor for AD ^12,13,36^. As modulation of PU.1 function is being investigated as a potential treatment for AD based upon results in model mice ^13^, we sought to explore the relationship between PU.1 and PPAR8. To do so, we evaluated our transcriptome results obtained from mouse brain microglia subjected to PPAR8 agonist treatment, and performed gene set enrichment analysis using the search engine Enrichr ^37,38^, selecting the ChIP-X Enrichment Analysis (ChEA) database to identify transcription factors whose binding sites are significantly over-represented in the microglia DEGs obtained upon PPAR8 agonist treatment (**Supplementary Table 5**). Interestingly, among the top hits from this analysis was PU.1. Of the 1,947 DEGs identified in microglia isolated from the brains of mice treated with PPAR8 agonist, ∼20% contain consensus binding sites for the transcription factor PU.1, suggesting that PPARδ and PU.1 are likely co-regulating a substantial number of genes in microglia.

Based upon these findings, we inquired whether PPARδ and PU.1 might physically interact by initially co-expressing PPARδ and PU.1 in HEK293T cells, and we observed evidence for an interaction (**Fig. 6a**). We then performed co-immunoprecipitation experiments for the endogenous proteins in iTF-microglia, and documented that PPARδ and PU.1 interact in these human microglia-like cells (**Fig. 6b**). To determine if PPAR8 agonism affects the interaction, we treated HEK293T cells with PPAR8 agonist, and noted that the interaction between PU.1 and PPARδ decreased in a dose-dependent manner (**Fig. 6c**). We then asked how PPAR8 agonism might affect PU.1 function, as reducing PU.1 activity has been demonstrated to be neuroprotective ^13^. To do so, we utilized an immortalized mouse microglia BV2 cell line stably transfected with a PU.1 luciferase reporter at the Rosa26 locus ^13^. When we treated this cell line with a PPAR8 agonist, we detected markedly reduced PU.1 transactivation (**Fig. 6d**). These findings suggest that one aspect of PPARδ regulation of microglia state may involve blunting PU.1 transcription activation.

**Figure 6.**
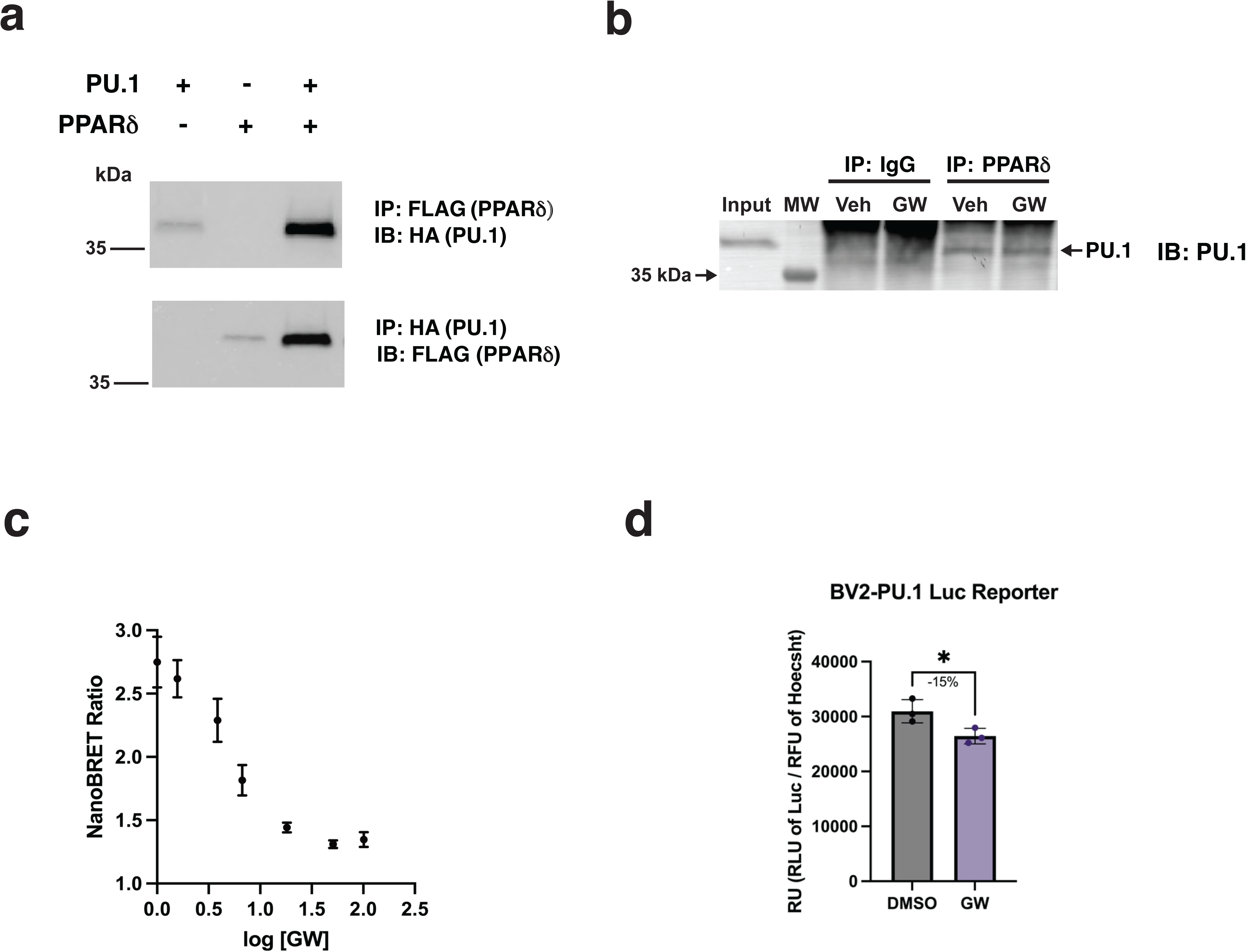
PPAR8 interacts with PU.1 and interferes with its activity. **a)** HEK293T cells were transfected with FLAG-tagged PPAR8 and 3xHA-tagged PU.1. Antibodies used for immunoprecipitation (IP) and immunoblot (IB) are as indicated. **b)** Co-immunoprecipitation in iTF-microglia with endogenous protein. Conditions as labelled. **c)** HEK293T cells were co-transfected with epitope-tagged PU.1 and PPAR8 constructs. Resultant fluorescence was measured and normalized to a NanoBRET ratio, with a higher ratio indicating greater interaction. n = 3 biological replicates. Error bars = s.e.em. **d)** BV2-PU.1-Luc cells were treated with or without GW501516 (1 μM), after which luminescence was measured. Luminescence values for each well were normalized to Hoechst signal. *p<0.05, two-tailed t-test, n = 3 biological replicates.

### PPAR8 attenuates inflammation in the CNS of Huntington’s disease and tauopathy mice

In light of our results indicating that PPARδ can suppress microglia inflammatory activation, and our previous findings showing that treatment of HD model mice with the brain-penetrant PPAR8 agonist KD3010 can dramatically rescue HD disease phenotpyes (17), we hypothesized that *in vivo* PPARδ neuroprotection might involve repression of neuroinflammation. To assess the status of inflammatory gene expression in the brains of the HD N171-82Q mice and test if PPAR8 agonist treatment affected neuroinflammation, we performed qRT-PCR analysis of NF-kB inflammatory pathway target genes, and detected significantly increased expression of these inflammatory targets in the striatum of HD mice (**Fig. 7a**). HD mice treated with KD3010, however, displayed a restoration of inflammatory gene expression back to control levels (**Fig. 7a**). We also assayed cytokine levels in plasma samples obtained from symptomatic HD mice, and in vehicle-treated HD mice, we noted a dramatic increase in TNF-α and IL-6 levels, which were markedly reduced in the plasma of HD mice treated with the PPAR8 agonist KD3010 (**Fig. 7b**). Levels of the anti-inflammatory cytokine IL-10 correspondingly increased in the plasma of KD3010-treated HD mice in comparison to vehicle-treated HD mice and non-transgenic controls (**Fig. 7b**). We then proceeded to test whether this finding could be extended to tau P301S (PS19) transgenic mice, an in vivo model of tau-mediated neurodegeneration. Increases in inflammatory genes were observed in the brains of PS19 mice treated with vehicle, but expression of these genes was markedly decreased in PS19 mice treated with KD3010 (**Fig. 7c**). To determine if this is a general effect of PPAR8 agonism, we also examined expression of these inflammation targets in wild-type C57BL/6J mice treated with the PPAR8 agonist KD3010, and documented repression of inflammation target genes in C57BL/6J mice treated with the PPAR8 agonist KD3010 (**Fig. 7d**). However, in mice expressing dominant-negative E411P PPARδ in the CNS, inflammation target gene expression was markedly increased (**Fig. 7e**). These findings indicate that PPAR8 activation is capable of blunting neuroinflammation *in vivo*.

**Figure 7.**
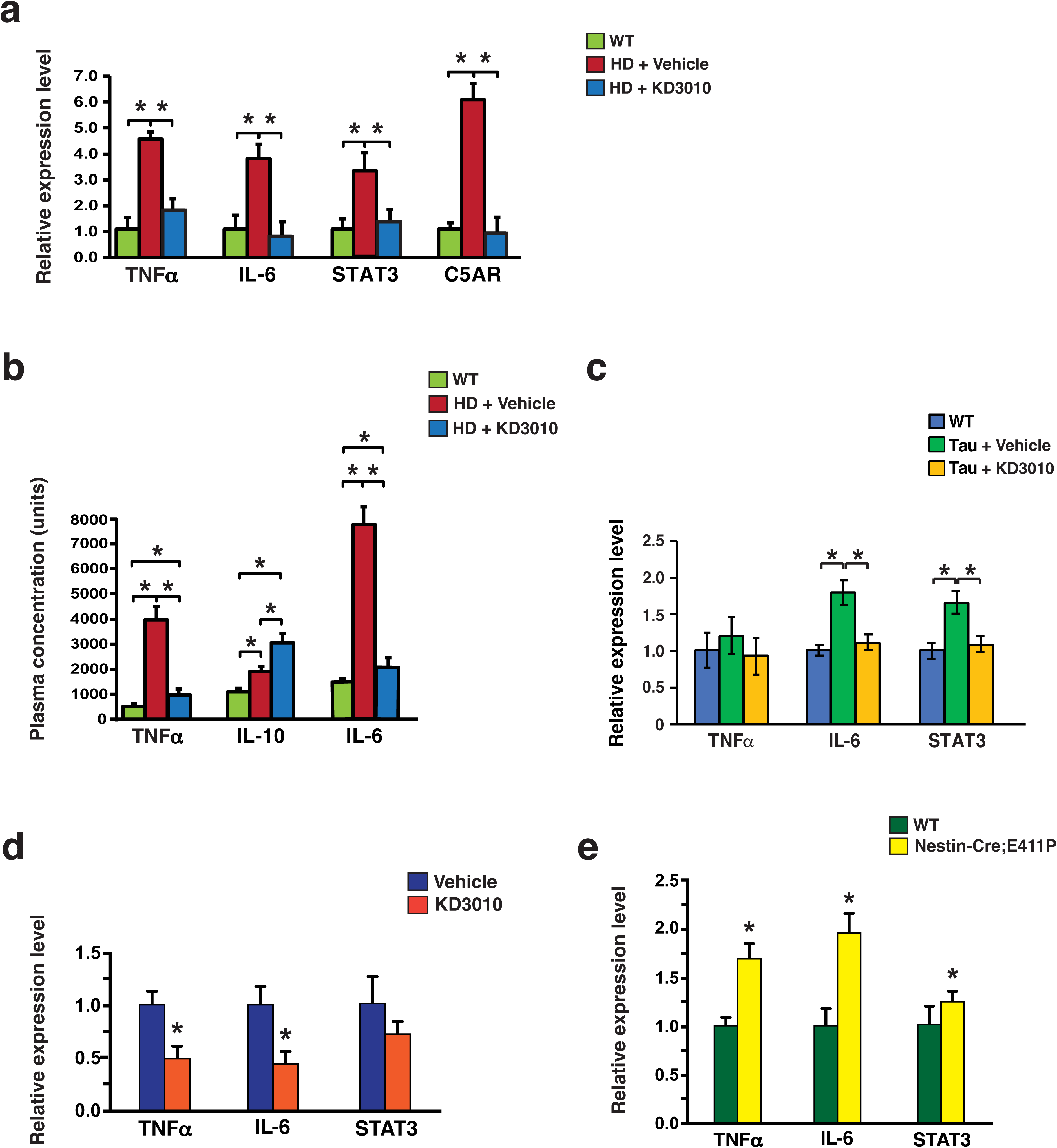
PPAR8 agonist treatment reduces inflammation in Huntington’s disease and tauopathy mice. **a)** HD N171-82Q mice were treated from 6 weeks of age with KD3010 (50 mg/kg/d) or vehicle. RNA was isolated from the striatum of 18-week-old HD and wild-type littermate control mice (n = 3 per group), and results of qRT-PCR amplification of inflammatory mediators are shown. *p < 0.05, ANOVA with post-hoc Dunnett test. **b)** Plasma from the same cohort shwn in ’A’was analyzed by flow cytometry for inflammatory mediators. *p<0.05, ANOVA with post-hoc Dunnett test. **c)** Tau P301S (PS19) mice were treated from 2 months of age with KD3010 (50 mg/kg/d) or vehicle. RNA from 6-month-old Tau P301S mice and wild-type littermate controls (n = 3 / group) was analyzed by qRT-PCR for inflammatory mediators. *p < 0.05, ANOVA with post-hoc Dunnett test. **d)** 10-week-old C57BL/6J wild-type mice were treated with KD3010 (50 mg/kg/d) or vehicle (n = 4 / group). Striatal RNA was analyzed by qRT-PCR analysis for inflammatory mediators are shown. *p<0.05, two-tailed t-test. **e)** RNA from the striatum of 8-month-old PPARδ-E411P; Nestin-Cre bigenic mice and wild-type littermates (n = 4–5 / group) was analyzed by qRT-PCR for inflammatory mediators. *p < 0.05, two-tailed t-test. Error bars = s.e.m.

## DISCUSSION

Understanding the role of microglia in neurodegenerative diseases such as AD has revealed the remarkable complexity of this cell type, and has underscored the need to characterize microglial state through a combination of strategies by evaluating function and transcriptome both *ex vivo* and *in vivo*. Because microglia are extremely dynamic, numerous studies have documented discrepant and opposing effects of microglia in the context of neurodegenerative disease, making their therapeutic modulation exceedingly difficult. Although depletion of microglia has been attempted in AD model mice, the results can differ based upon disease stage ^39^. For these reasons, identification and characterization of master transcription factors that can modulate microglia activity and function will be crucial for advancing therapy development to treat AD and related disorders. In this study, we present a series of findings that indicate how and why PPARδ deserves consideration as a microglia drug target. Our results indicate that PPAR8 agonism promotes a microglial state that upregulates phagocytosis and lipid processing, while reducing inflammation. Endocytic and lipid processing pathways were upregulated by PPAR8 agonism in single-cell transcriptome analysis of iTF-microglia, and PPAR8 agonism correspondingly increased the phagocytosis of Aβ and rat synaptosomes in iTF-microglia. Supporting this, PPARδ was identified as a positive hit in an unbiased CRISPRa screen for modulators of phagocytosis in iTF-microglia ^22^.

One useful approach for characterizing microglia cell state in iMGs in a way that mimics a neurodegenerative disease insult is by exposing iMGs to materials from neuron andn other CNS cellular debris, using a formulation known as “brain powder” ^33^. Here we tested the effect of both a PPAR8 agonist and a PPAR8 antagonist in iTF-microglia exposed to brain powder, and we monitored the shift in six distinct microglia transcriptome states. The most striking aspect of this analysis was the diametrically opposite effect on the Mg3 cluster, which was most highly enriched for inflammatory response genes, as PPAR8 agonism reduced the Mg3 cluster, while PPAR8 antagonism increased the Mg3 cluster. These findings corroborated our transcriptome results obtained in microglia isolated from the brains of mice treated with the PPAR8 agonist KD3010. Another defining feature of PPAR8 agonism in microglia is a shift out of a homeostatic state to a primed DAM-like state, which was reflected by the ability of PPAR8 activation to repress the expression of *P2RY12*, and promote the expression of endocytic and lipid processing genes, such as *ABCA1* and *PLIN2*. However, PPAR8 agonism did not perfectly recapitulate all the gene expression changes that comprise the core DAM transcriptome signature, indicating that the effect of PPAR8 is unique and distinct, consistent with an emerging view that the DAM state is not monolithic, but rather dynamic based upon context ^3^. This nuanced effect of PPAR8 activation on microglia was also captured in a recent large study of 194,000 single-nucleus microglial transcriptomes and epigenomes isolated from 217 AD patients and 226 AD controls, where PPARδ expression was found to be significantly increased in the MG2 inflammatory I cluster in AD patients ^6^. The MG2 inflammatory I cluster was described as an early stage along the progression to frank inflammation, as cytokine signaling pathway, cytokine, and NF-kb signaling pathway genes were yet to be highly expressed, but the MG2 cluster clearly represented a strong shift away from the homeostatic state. Of note, snATAC-seq analysis of 41,832 microglia nuclei also identified the PPARδ promoter as a significantly enriched peak in AD patient microglia ^6^, suggesting that altered PPARδ regulation occurs in AD. Another interesting result from this combined transcriptome-epigenome analysis of AD patient microglia was the detection of increased levels of PPARγ with increased expression of *P2RY12* in the brains of AD patients, where PPARγ expression was noted to be highest in the MG4 lipid processing cluster and was often accompanied by increased expression of *APOE*, yielding an atypical DAM-like state ^6^. Hence, while PPAR8 and PPARγ are both highly expressed in microglia, and both can promote phagocytosis in microglia based upon our results here and previously published work on PPARy^40^, over-expression of PPARγ in microglia has recently been shown to maintain microglia in a homeostatic state ^6^, whereas PPAR8 is likely favoring a shift away from compensatory homeostasis and toward a more activated phagocytosis-primed state with an inflammatory tendency, but without frank inflammation.

As previous studies have shown that PPAR8 is capable of transrepression of gene expression (23, 24), we considered the possibility that PPARδ might somehow be opposing the effects of PU.1, a well-known microglia transcription factor whose gene targets strongly promote inflammation. PU.1 has been repeatedly identified as an AD risk gene, with increasing expression correlating with greater susceptibility to disease ^12,13,36^, and can prevent *APOE4*-expressing microglia from taking on a DAM-like state ^10^. Furthermore, repression of PU.1 in AD model mice increased cognition while decreasing neuroinflammation and neuronal loss ^13^, underscoring the importance of identifying factors that antagonize PU.1 activity. The PPARs and PU.1 have not been previously studied together in microglia; however, a functional interaction between PPARγ and PU.1 was shown in adipocytes, where PU.1 overexpression altered the cistrome of PPARγ, though no physical interaction was reported ^41^. We found initial evidence for a physical interaction between PPARδ and PU.1, and also noted that PPAR8 agonism could reduce PU.1 transcriptional activity, albeit in a transformed microglia cell line. Although further work will need to be done to fully elucidate the regulatory relationship between PPARδ and PU.1, our results suggest that PPARδ could exert neuroprotection in AD by blunting the effects of PU.1 in microglia. Because the PPAR8 agonist T3D-959 is moving forward in human clinical trials as a potential treatment for AD ^19^, delineating whether PPAR8 is capable of transrepression of PU.1 could provide further insight into its mechanism of action in AD. It is also possible that transrepression of PPARδ by PU.1 may contribute to the ability of PU.1 to prevent microglia from transitioning to a DAM-like state, since we found that PPAR8 inhibition blunted expression of the transcription factor MITF, which has now been implicated as a driver of DAM gene expression and phagocytic activity ^33^.

Given the interest in PPAR8 agonist therapy as a treatment for neurodegenerative disease in human patients and its recent clinical trial experience in AD, we chose to pursue two small pilot studies in model mice to evaluate if PPAR8 activation could blunt neuroinflammation *in vivo*. Because we have already shown that the PPAR8 agonist KD3010 is an effective treatment in HD N171-82Q mice (17), we first pursued studies in these HD mice, and after noting increases in NF-kB dependent inflammatory mediators, we tested if KD3010 could reduce such neuroinflammation. In the CNS of KD3010-treated HD mice, we observed a marked decrease of inflammatory mediators down to levels seen in non-disease wild-type littermate control mice. This was paralleled by marked reductions in inflammatory mediators in peripheral blood plasma. We then selected an AD-relevant model – the tau P301S (PS19) tauopathy mice ^42^, and evaluated the effect of KD3010 treatment, again documenting significant decreases in levels of inflammatory mediators to roughly normal, confirming that PPAR8 agonism can modulate neuroinflammation. These findings corroborate our transcriptome results in iTF-microglia, where PPAR8 agonism blunted inflammatory gene expression associated with the transcription factors NFKB1, RELA, and PU.1, and are consistent with analysis of microglia transcriptomes obtained from KD3010-treated wild-type mice. Indeed, the ability of PPARδ to reduce neuroinflammation was observed in wild-type mice subjected to KD3010 agonist treatment, and loss of PPARδ function via expression of a dominant-negative mutant version of PPARδ alone was sufficient to induce neuroinflammation. Favorable effects of PPAR8 activation have also been demonstrated in a mouse model of EAE, where PPAR8 agonism reduced inflammation and was neuroprotective ^21^, indicating that PPAR8 agonist therapy might be beneficial in a variety of neurological diseases.

Although our findings across multiple model systems both *ex vivo* and *in vivo* indicate that PPAR8 activation may achieve neuroprotection by preventing excessive neuroinflammation, many open questions remain and there are limitations to our results. One concern is that cell autonomous effects of PPAR8 in iTF-microglia do not precisely mimic microglial behavior *in vivo*. Furthermore, in the brains of HD and tauopathy model mice, microglia are surrounded by various other CNS cell types; hence, microglia may not be the only source of measured inflammatory mediators. Finally, our initial evaluation of the biochemical interaction between PPARδ and PU.1 was restricted to transformed cell lines, and will require further examination in microglia *in vivo*. In addition to these caveats, understanding the role of PPARδ in regulating lipid metabolism in microglia in different disease contexts will be an important next step, as lipid metabolism is closely tied to inflammation ^6,43–45^, and aberrant lipid droplet accumulation in microglia, which is promoted by *APOE4* expression, can result in excessive reactive oxygen species and secretion of proinflammatory cytokines ^11^, underscoring the importance of this phenomenon in AD. Despite these unanswered questions, our results indicate that PPAR8 is a master transcription factor in microglia, sitting at the nexus of key pathways that dictate microglia state and function. Therefore, pharmacological modulation of PPAR8 may represent an appealing therapeutic opportunity for many neurodegenerative disorders, including AD.

## METHODS

### Mouse Treatment Protocol

Eight wild-type (C57BL/6J Jax Strain #:000664) mice between the ages of 6 and 8 months were divided into two cohorts of 4 mice each. One group received intraperitoneal (IP) injections of KD3010 5 days per week at a dose of 50 mg/kg while the other received vehicle IP injections of the same frequency. Injections were carried out for a total of six weeks. KD3010 was prepared at a concentration of 10 mg/mL in Corn Oil with 5% DMSO and vortexed immediately prior to administration.

### Microglia Isolation

Microglia were isolated from the brains of mice using a previously established protocol ^46^. Briefly, whole brains from mice were first finely chopped, then dissociated by trituration with Pasteur pipettes in a dissociation solution containing Collagenase A and DNase I. Microglia were then isolated using CD11b microbeads and a magnetic stand. Samples from the whole tissue lysate before isolation and flowthrough samples after isolation were harvested for further analysis. Following subsequent washes, isolated microglia and other samples are stored at -80°C in Trizol for RNA processing.

### Human iPSC Culture

Human iPSCs were cultured in mTeSR Plus Medium on 60 mm TC-treated Culture dishes coated with LDEV-free Corning Matrigel hESC-Qualified Matrix diluted 1:100 in Dulbecco’s Modified Eagle’s Medium (DMEM), 1X. Media was replaced daily. Cells were ReLeSR passaged when 70-80% confluent. For experiments, cells were dissociated with Accutase, counted using the Countess 3 Automated Cell Counter and plated with Y-27632 dihydrochloride ROCK inhibitor diluted 1:1000 in differentiation media.

iTF-iPSCs on a KOLF2.1J background were graciously provided to us by the lab of Dr. Bill Skarnes. To freeze iTF-iPSCs for future use, old media from plates was aspirated. Then, the plate was rinsed with 1x DPBS and treated with ReLeSR. ReLeSR was aspirated within 1 minute, and plates were incubated at 37^°^C for 10 minutes. Plates were then treated with DMEM, 1X with strong enough force to dislodge iTF-iPSCs. Free floating cells were then transferred to a 15mL conical tube and centrifuged at 50xg for 3 minutes. The resulting supernatant was aspirated, and cells were resuspended in 1mL of CryoStor CS10 before being transferred to CryoPure 1.6mL tubes and placed into a Mr. Frosty Freezing Container at -80^°^C overnight. No later than 48 hours later, the cells were transferred to -160^°^C for long term storage.

iTF-iPSCs that are stored as frozen aliquots were thawed in a 37^°^C water bath, and diluted 1:15 in mTeSR Basal Medium. Diluted iTF-iPSCs were transferred to a 15mL conical tube and centrifuged at 220xg for 5 minutes before aspirating the supernatant. Cells were then resuspended in mTeSR Plus Basal Medium supplemented with 10nM Y-27632 dihydrochloride ROCK inhibitor and plated onto Matrigel-coated plates. After one day, media was aspirated and replaced with mTeSR Plus Basal Medium.

### Differentiation of iTF-Microglia

iTF-iPSCs were differentiated using a previously established protocol^22^. Cells were cultured for at least 24 hours without Y-27632 dihydrochloride ROCK inhibitor before being dissociated with Accutase. Cells were incubated at 37°C for 7 minutes with Accutase, after which time the Accutase was quenched by adding DMEM at a 3:1 ratio. Dissociated cells were transferred to a 15mL conical tube and centrifuged at 220xg for 5 minutes. After aspirating the resulting supernatant, cells were resuspended in Day 0 Differentiation Media: Essential 8 Basal Medium containing 10nM ROCK inhibitor and 2 μg/ml Doxycycline hyclate. iTF-iPSCs were counted and plated (Day 0) onto plates double-coated with Poly-D-Lysine Coated and Matrigel. After two days (Day 2), the media was aspirated and changed to Day 2 Media: Advanced DMEM/F12 Medium containing 2 μg/ml doxycycline, 1x GlutaMAX^TM^ Supplement, 100 ng/ml Human IL-34, and 10 ng/ml Human GM-CSF. After two additional days (Day 4), the media was aspirated and replaced with Day 4 media: Advanced DMEM/F12 containing 1X GlutaMAX, 2 μg/ml doxycycline, 100 ng/ml Human IL-34 and 10 ng/ml Human GM-CSF, 50 ng/ml Human M-CSF and 50 ng/ml Human TGF-β1. Four days later (Day 8), cells were fully differentiated (termed iTF-microglia) and ready for use. Media was replaced on Day 8 and every 3-4 days as needed with the same Day 4 media. Differentiation was confirmed by morphology and immunocytochemistry.

### LPS Treatment/Cytokine Array

iTF-Microglia were treated with 100 ng/mL LPS diluted in DPBS, as done previously^22^, with or without the addition of PPAR8 agonism. The inflammatory readout was assessed by analyzing inflammatory cytokine production using an Abcam Human Cytokine Array for 42 targets. After 24 hours of treatment with or without LPS and with or without 1 μM GW501516, the conditioned media from the iTF-Microglia was collected and added to the cytokine array per the manufacturer’s instructions. After taking chemiluminescence images of the blots using a Bio-Rad ChemiDoc Imaging System, the images were analyzed using the Protein Array Analyzer tool on FIJI to quantify the amount of protein.

### Phagocytosis Assay

To assess phagocytic activity of iMGLs and the ability of a PPAR8 agonist to alter it, we utilized fluorescent fibrillar beta-amyloid (1-42) fAβ_1-42_ and pHrodo tagged rat synaptosomes. To prepare fluorescently labeled fAβ_1-42_, we followed a previously published protocol ^47^. The lyophilized stock was first dissolved in NH_4_OH to an initial concentration of 1 mg/mL, then further diluted to 100 μg/mL with UltraPure^TM^ DNAse/RNAse free water and incubated for 7 days at 37°C. To isolate rat synaptosomes, we obtained Innovative Grade US Origin Rat Sprague Dawley Brain. Then, following a previously reported protocol ^22^, 10 mL / gram of brain of Syn-PER Synaptic Protein Extraction Reagent was supplemented with protease and phosphatase inhibitor was added to the brain and was used to homogenize using a Dounce homogenizer on ice. The contents were then put into a tube and were centrifuged at 1200xg for 10 minutes at 4°C. Then, the pellet was discarded and this centrifugation was repeated using a new tube. After that repeat spin-down, the supernatant was spun down at 15,000xg for 20 minutes at 4°C. After discarding the supernatant, the wet pellet was weighed and the synaptosome was resuspended at a concentration of 50 mg/mL. Then, in order to conjugate these to the pH-sensitive pHrodo dye, 3μM of pHrodo-Red was added to the synaptosomes and this mixture was incubated for 45 minutes in the dark at room temperature. Then, the synaptosomes were diluted 1:10 in DPBS and spun down at 2500xg for 5 minutes. After two more washes, the synaptosomes were resuspended at 50mg/mL in Advanced DMEM/F12 with 5% DMSO for future use.

Day 9 iTF-Microglia were used for phagocytosis assays. We had conditions where iPSC cultures are combined with 2 μg/mL fluorescent fibrillar beta-amyloid or 50 μg/mL pHrodo tagged human synaptosomes, with or without 1μM GW treatment, as well as a no substrate condition. We incubated these cells for 24 hours in an Incucyte S3 at 37°C to allow for phagocytosis. We analyzed using the Incucyte S3 software and normalized to fluorescence intensity at t=0.

### Migration Assay

To measure migration of iTF-Microglia, 20,000 iTF-Microglia were plated onto PDL and ECM coated 8 μm transwells in assay media with GW, DMSO, or DMSO/ADP. The transwells were placed in 24-well plates containing assay media (Advanced DMEM/F12 + 1X GlutaMAX + 2 μg/mL Doxycycline) and GW or DMSO. The cells were incubated for an hour at 37°C. Attractants (5 μg/mL fAβ_1-42_, 5 μg/mL P301S Tau preformed fibrils, 1 mg/mL synaptosomes, 35,000 apoptotic SH-SY5Y / cm^2^, and 100μM ADP were added to the bottom of 24-well plates and the transwells were moved onto them. Fibrillar Aβ and synaptosomes were prepared using a standard protocol ^33^. The cells were incubated with the attractants for 4 hours at 37°C. The cells were then fixed with 4% PFA and stained with Hoechst. The cells on the top of the transwells were wiped away and the remaining migrated cells on the bottom of the transwells were imaged using an Echo Revolution microscope and the total migrated cells were compared.

### RNA Extraction and qRT-PCR

RNA was extracted using Qiagen RNeasy Mini RNA extraction kit, converted to cDNA with SuperScript IV VILO Master Mix with ezDNase Enzyme, and genes of interest were tested for expression change using qRT-PCR. Both TaqMan assays and SyBr green assays were utilized in this paper depending on the target. Relative gene expression using *Tfrc and Actb* as normalization controls was calculated using 2^-ddCt^. Primer sequences are as follows:

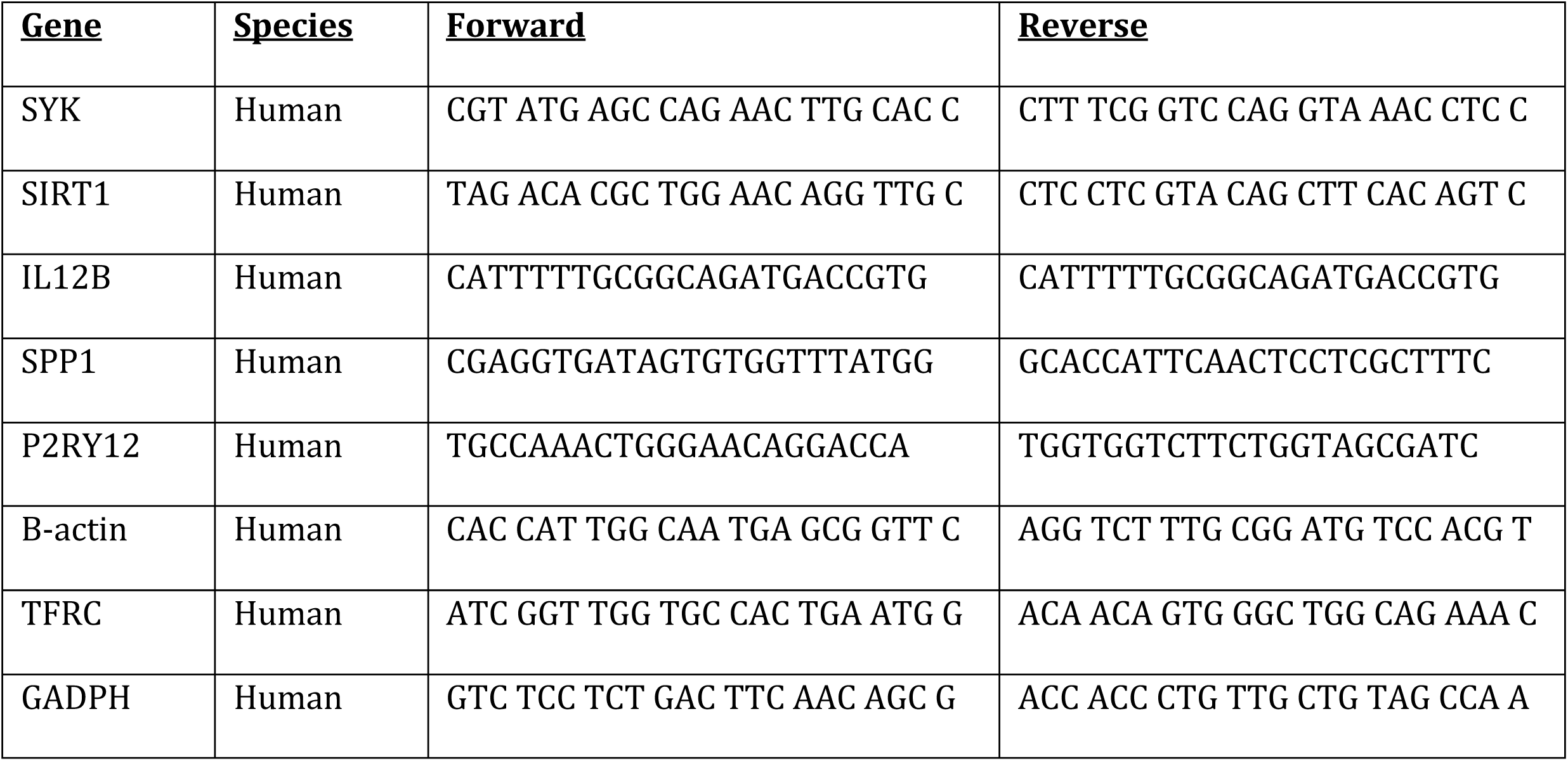

### Preparation of Mouse Brain Powder

Twelve-month old female wild-type mice were perfused with ice-cold PBS and their brains were isolated. Olfactory bulbs and cerebellum were removed from the brain, and the brain was then cut in half and immediately flash frozen with liquid nitrogen. Frozen brains were made into powder using a tissue pulverizer, all while kept frozen, and were weighed out to predetermined amounts based on experimental needs. On the morning of the experiment, the mouse brain powder was resuspended at a concentration of 3.33 mg / 100 μL of sterile 1X DPBS. The powder was then homogenized by using a handheld Eppendorf tube homogenizer 10 times. Then, the powder was tun through a 70 μm filter to remove any remaining large clumps.

### Treatment of iTF-Microglia with Brain Powder

Day 8 iTF microglia were plated in 24 well plates at a density of 100,000 cells/well. 20 μL of brain powder solution was added to 36 wells and 20 μL of 1X DPBS was added to an additional 36 wells. iTF-Microglia were then treated with vehicle, 1 μM GW501516 or 1 μM GSK3787 (12 wells per treatment for both DPBS and brain powder groups). After 24 hours iTF-Microglia were collected and underwent a cleanup protocol to remove residual brain powder debris. Briefly, cells were lifted using TrypLE as described above, three wells were pooled for each replicate (n=4 replicates per treatment group), and cells were spun at 200xg for 5 minutes. Supernatants were aspirated, each pellet was resuspended in 1 mL of Advanced DMEM/F12, and each replicate was run through a 70 μM filter into a new 15 mL tube. An additional 5.2 mL of Advanced DMEM/F12 was added to the cells and the tube was mixed. 1.8 mL of standard isotonic Percoll (SIP; 9 parts Percoll + 1 part 10X PBS) was added to each sample and the tube was mixed by inversion three times. Then, 2 mL of 1X DPBS was gently added on top of the SIP/cell mixture. The tubes were then centrifuged at 400xg for 20 minutes with acceleration at 0 and brake at 4. Supernatants were aspirated, pellets were resuspended in 1mL of 0.5% BSA in 1X DPBS, followed by an additional 2 mL of 0.5% BSA. The samples were centrifuged again at 200xg for 5 minutes, supernatants were aspirated, and pellets were resuspended in 105 μL of the 0.5% BSA solution. Cells were then fixed using the Parse Biosciences Evercode^TM^ Cell Fixation v2 kit according to the manufacturer’s protocols.

### Parse scRNA-seq Barcoding, Sequencing, and Alignment

Fixed cell samples were barcoded using the Evercode^TM^ WT v2 kit according to manufacturer’s procedures. In short, samples were evenly loaded at a concentration aimed to capture 100,000 total cells. Once the initial barcoding was completed, pooled samples were split into 8 identical sublibraries containing 12,500 cells each. Illumina libraries were generated according to the Parse guidelines and each sublibrary was sequenced at a depth targeting ∼50,000 reads per cell. FASTQ files were aligned to the Ensembl GRCh38 transcriptome (filtered to keep protein coding genes) using the Parse split-pipe v1.1.2 alignment program with default settings. In total, 74,014 cells were identified across all eight sublibraries with an average read depth of 63,370 reads per cell, resulting in an average of 3,084 cells per sample and an average of 12,335 cells per experimental group.

### scRNA-seq Quality Control

Samples were then imported into Seurat v5 ^48^ for quality control preprocessing. Initially, each sublibrary was cleaned individually by normalizing, scaling, and clustering data using default settings and parameters. Doublets were identified using scDblFInder ^49^, and removed as were clusters displaying high levels of cell division genes (**Supplementary Table 2**). Remaining cells were then reclustered and clusters with low gene counts or residual clusters with high cell division scores were removed. Finally, remaining cells were filtered to exclude all cells with mitochondrial read percentage greater than 1.1%, ribosomal read percentage greater than 1.5%, UMI and gene counts less than 2,000 and greater than twice the median count.

Following the individual sublibrary QC, all eight sublibraries were merged into a single Seurat object, data was normalized, scaled while regressing out the ‘Sublibrary’ variable, and reclustered. Clusters were identified that expressed high levels of non-microglial genes (incompletely differentiated cells) and high levels of genes associated with p53 signaling (**Supplementary Table 2**). After removal of these clusters, a total of 33,745 cells remained with an average of 3,888 genes and 9,733 UMI per cell.

### scRNA-seq Data Integration, Differential Gene Analysis, and Visulization

#### Response to PPARδ Agonism and Antagonism

All saline treated samples (no brain powder) were extracted from the combined Seurat object, normalized using default settings, and scaled using the top 2,000 variable features (followed by removal of mitochondrial and ribosomal genes) while regressing out the ‘Sublibrary’, ‘nCount_RNA’, and ‘percent.ribo’ variables. Individual layers were then integrated using the ‘IntegrateLayers’ function and the “Harmony” integration method. Integrated samples were clustered using the first 50 PCs and a resolution of 0.5. Data layers were joined using the ‘JoinLayers’ function and pseudobulk normalized gene counts were generated using the ‘AggregateExpression’ function to group the data by replicate and treatment group. Differential gene expression analysis was then performed on the aggregated expression data using the ‘FindMarkers’ function to perform pairwise comparisons between treatment groups. Box plots were generated in Graphpad Prism v10.4.1 using the aggregated expression data. All genes presented reached significance in at least one pairwise comparison in Seurat and presented data for all three groups was reanalyzed in Prism using a one-way ANOVA with Tukey’s test for multiple comparisons.

#### Response to Brain Powder Treatment

All vehicle treated samples (no PPARD modulation) were extracted from the combined Seurat object, normalized using default settings, and scaled using the top 2,000 variable features (followed by removal of mitochondrial and ribosomal genes) while regressing out the ‘Sublibrary’, ‘nCount_RNA’, and ‘percent.ribo’ variables. Individual layers were then integrated using the ‘IntegrateLayers’ function and the “Harmony” integration method. Integrated samples were clustered using the first 28 PCs and a resolution of 0.55. Data layers were joined using the ‘JoinLayers’ function and pseudobulk normalized gene counts were generated using the ‘AggregateExpression’ function to group the data by replicate and treatment group. The DAM signature was calculated for all cells using a list of genes that has been consistently upregulated in our previously published scRNA-seq data ^50–52^ and the ‘AddModuleScore’ function. Differential gene expression analysis was then performed on the aggregated expression data using the ‘FindMarkers’ function to perform pairwise comparisons between treatment groups. The violin and box plots were generated in Graphpad Prism v10.4.1 using the module score or aggregated expression data, respectively. All boxplot genes presented reached significance in Seurat, but presented data for all plots displays significance levels from two-tailed t tests. Pathway and transcription factor prediction analyses were performed using Metascape’s express analysis^34^ with genes that were significantly upregulated following powder treatment (FDR<0.05; LFC>0.25) used as input. All analysis parameters and data are shown in **Supplementary Table 3**.

#### Effect of PPARδ Modulation on Powder Response

All brain powder treated samples (no saline treated samples) were extracted from the combined Seurat object, normalized using default settings, and scaled using the top 2,000 variable features (followed by removal of mitochondrial and ribosomal genes) while regressing out the ‘Sublibrary’, ‘nCount_RNA’, and ‘percent.ribo’ variables. Individual layers were then integrated using the ‘IntegrateLayers’ function and the “Harmony” integration method. Integrated samples were clustered using the first 50 PCs and a resolution of 0.5. Data layers were joined using the ‘JoinLayers’ function and pseudobulk normalized gene counts were generated using the ‘AggregateExpression’ function to group the data by replicate and treatment group. Pathway and transcription factor prediction analyses were performed using Metascape’s express analysis.

Differential gene expression analysis was performed between the GW501516-treated and GSK3787-treated aggregated data samples and the upregulated genes (FDR<0.05; LFC>0.25) or downregulated genes (FDR<0.05; LFC<-0.25) were used as the input for PPARD agonism and PPARD antagonism analyses, respectively. The ‘Inflammatory Response’ and ‘Metabolism of Lipids’ signatures were calculated for all cells using a subset of genes from the GO Biological processes ‘Inflammatory Response’ pathway or the Reactome ‘Metabolism of Lipids’ pathway (**Supplementary Table 4**) and the ‘AddModuleScore’ function. Differential gene expression analysis was then performed on the aggregated expression data using the ‘FindMarkers’ function to perform pairwise comparisons between treatment groups. Box plots were generated in Graphpad Prism v10.4.1 using the aggregated expression data. All boxplot genes presented reached significance in at least one Seurat pairwise comparison and/or as cluster-specific genes, but presented data for all three groups was reanalyzed in Prism using a one-way ANOVA with Tukey’s test for multiple comparisons. All analysis parameters and data can be found in **Supplementary Table 4**.

#### Transfection

HEK293T cells were transfected with plasmids utilizing Lipofectamine 3000 according to manufacturer’s instructions. Cells were cultured in 10cm plates and transfected with 14μg total of plasmid for 24 hours before media was replaced and treatment was administered.

#### Co-immunoprecipitation

Cells from different treatment groups were lysed using IP lysis buffer. After lysis, the lysate was spun at 13,000xg for 15 minutes and the concentration of the lysate was calculated by BCA on the supernatant of the spin-down. At this point, the samples were further diluted to a final concentration of 1 μg/mL. For immunoprecipitation, samples were incubated overnight at 4**°**C with anti-FLAG beads or 5 μg antibody according to manufacturer’s instructions. If just incubated with the antibody, the next day, Protein A beads were added and the samples were incubated on a rotator for 1.5 hours at 4**°**C . After the incubation, the flow-through was saved and the antigen-antibody-bead complex was transferred to a new tube for elution after a series of washes with IP lysis buffer and water. The samples were eluted by heating samples at 70°C for 10 minutes in a solution composed of IP lysis buffer, 4x LDS Sample Buffer and 10x Reducing Agent. After this, the sample was removed from beads using a magnetic rack and samples were loaded into a gel for western blot analysis.

#### NanoBRET

HaloTag and NanoLuc tagged plasmids for PPARδ and PU.1 were prepared using a NanoBRET starter kit from Promega. All combinations of N- vs C- tagged plasmids were tested and the best combination providing dynamic range was utilized. HEK293T cells were plated at a density of 350,000 cells in a 6-well plate and allowed to settle overnight. The following day, cells were transfected with appropriate plasmids using Lipofectamine 3000, with p53 and MDM2 being used as a positive protein interactor control, using manufacturer’s recommendations. The following day, cells were replated in a 96 well plate and treated with predetermined concentrations of GW501516, with concentrations ranging from 1 nM to 10 μM. The following day, the NanoBRET assay was carried out in the 96-well plate and absorbance readings were taken at 460nm (donor emission) and 618nm (acceptor emission), with a NanoBRET ratio being calculated after removing background absorbance.

#### BV2 PU.1 Lucerase Assay

BV2 cells with the PU.1 luciferase reporter were graciously provided to us by the lab of Dr. Li Huei Tsai. Briefly, these cells contain five tandem copies of the lB motif with a promoter and luciferase coding sequence from the pGL4.23 Promega luciferase plasmid inserted into the ROSA26 locus of the cells^13^. To do this, 10,000 cells were plated onto 96-well plates (Nunc 136101 for luciferase, Cellvis P96-1.5P for Hoechst) and treated with 1µM GW or vehicle. Cells were incubated for 48 hours with media change twice a day. Luciferase activity of the replicates was tested using ONE-Glo™ EX Luciferase Assay System (Promega A8110) according to the manufacturer’s instructions on a Thermo Scientific VarioskanLux. Cells were then stained with Hoechst at 1:10,000 for 15 minutes for normalization. The VarioskanLux was subsequently used to measure the Hoechst signal (Excitation 350 nm, Emission 461 nm).

## ACKNOWLEDGEMENTS

iTF-iPSCs on the KOLF2.1J background were generated and generously provided by Dr. Bill Skarnes. BV2-PU.1-Luciferase cells were graciously provided by the lab of Dr. Li Huei Tsai. Plasmids for NanoBRET experimentation were kindly generated by Dr. Bryce L. Sopher. This work was supported by grants from the N.I.H. (F30AG081084 to J.S.D.; R35 NS122140 and RF1 AG033082 to A.R.L.S.)

## Availability of Data

All processed data and raw sequencing reads will be made available (accession numbers pending).

## Author Contributions

J.S.D., J.H., M.B.-J., and A.R.L.S., provided the conceptual framework for the study. J.S.D., J.H., M.B.-J., and A.R.L.S. designed the experiments. J.S.D., J.H., U.R., Y.G., L.S., B.C., C.K., D.T., M.-H.V.T., A.G., and A.S.D. performed the experiments. J.S.D., J.H., and A.G. analyzed the data. J.S.D., J.H., and A.R.L.S. wrote the manuscript..

## Supplementary Figure Legends

**Figure 1.**
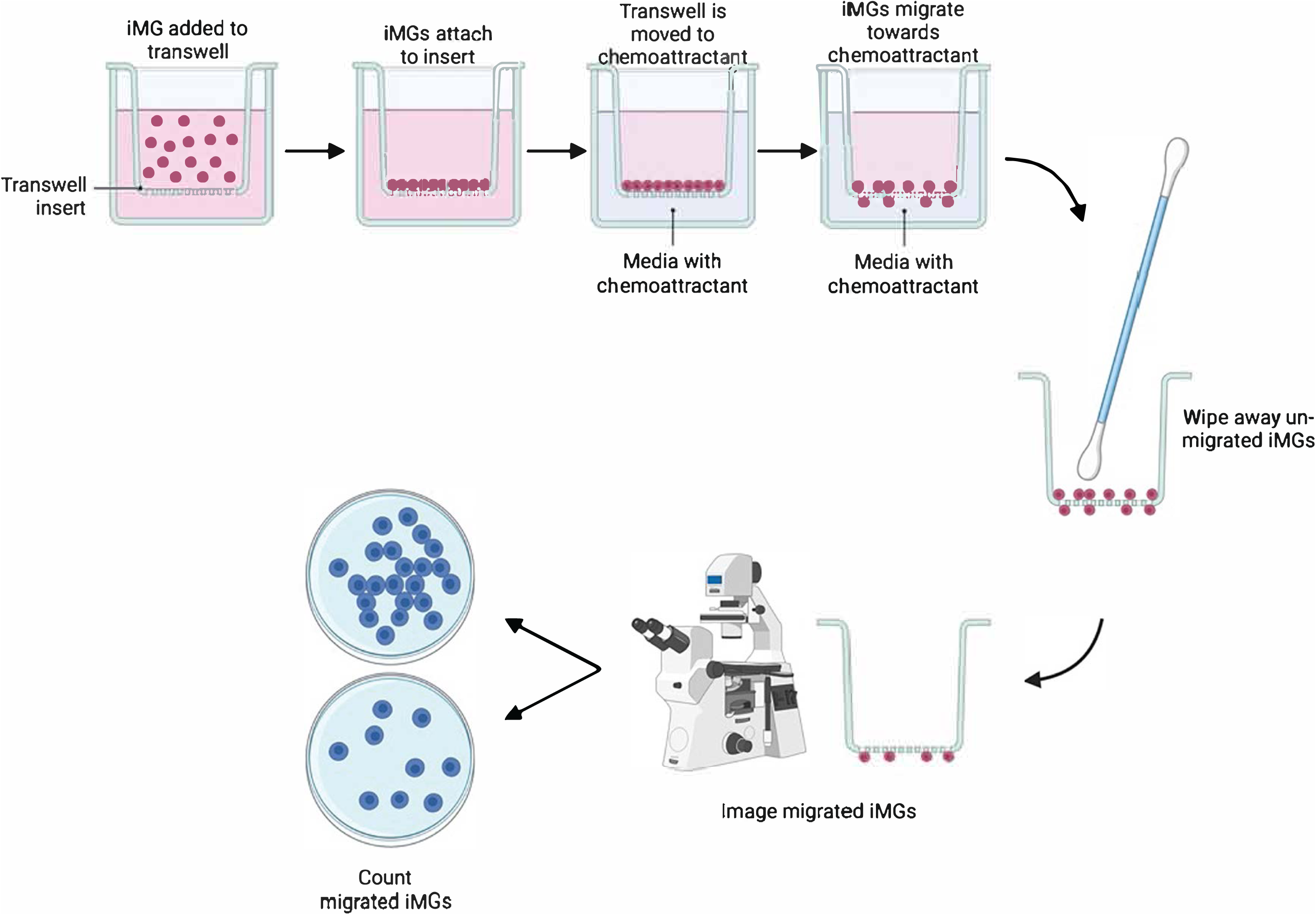
Set-up for migration assay protocol. Here we see a step-by-step diagram of the approach used to monitor microglia migration in response to a specified chemoattractant.

**Figure 2.**
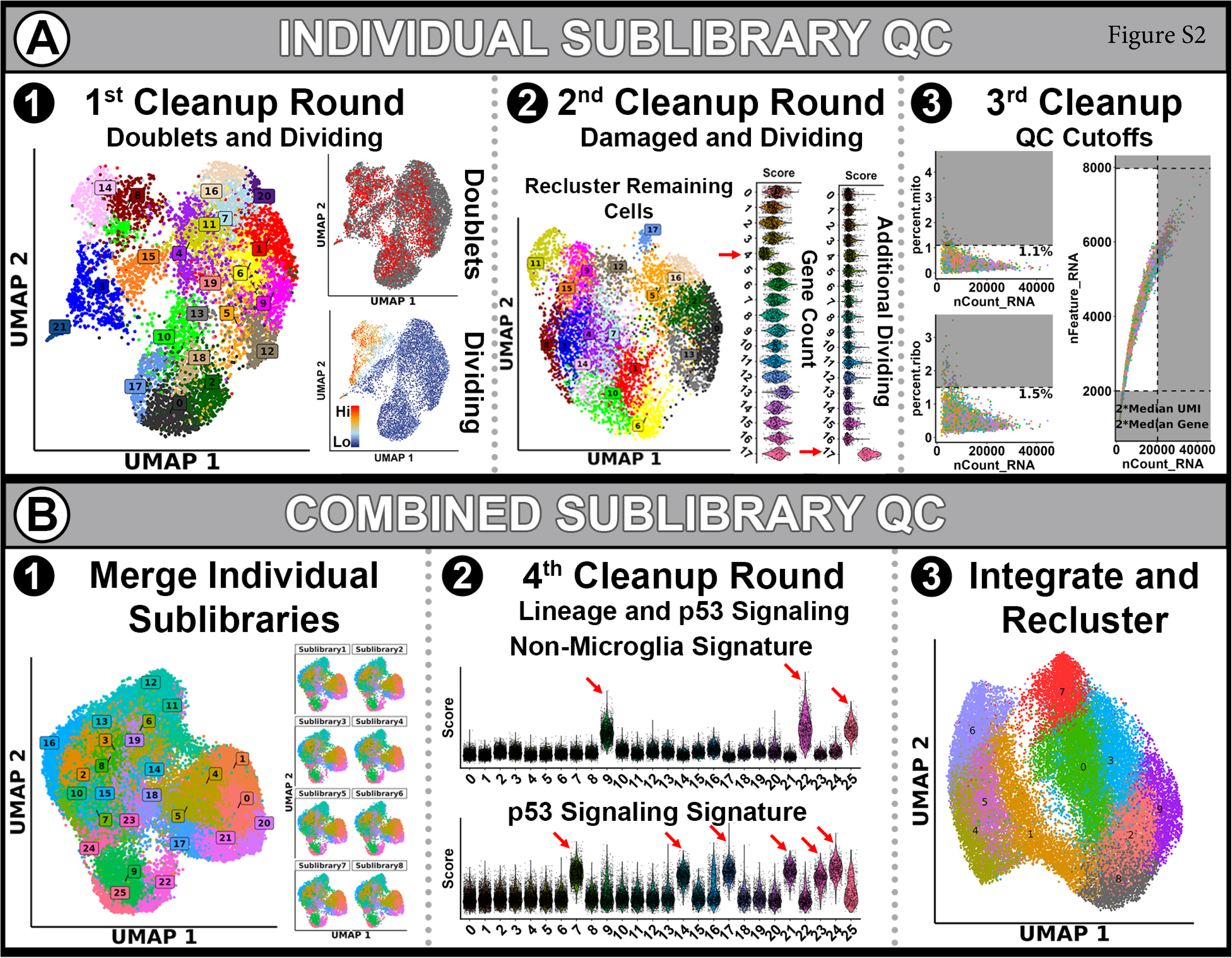
Quality control steps for Parse single cell sequencing. **a)** Each sublibrary underwent quality control clean-up individually prior to object merging. Cells were clustered followed by removal of doublets and dividing cells (1). The remaining cells were reclustered and cells with low gene counts and residual dividing cells were removed (2). Finally, cutoffs were applied to the remaining cells to remove cells with greater than 1.1% mitochondrial reads, greater than 1.5% ribosomal reads, cells with less than 2000 genes, and cells with greater than double the median number of genes or UMIs (3). **b)** Once each sublibrary underwent quality filtering, sublibraries were combined and reclustered as a single object (1). Additional clusters were removed that were found to express high levels of non-microglial gene signatures or high levels of genes related to p53 signaling (2). The remaining cells were then integrated prior to downstream analysis (3).

